# The salt-and-pepper pattern in mouse blastocysts is compatible with signalling beyond the nearest neighbours

**DOI:** 10.1101/2023.05.04.539359

**Authors:** Sabine C. Fischer, Simon Schardt, Joaquín Lilao-Garzón, Silvia Muñoz Descalzo

## Abstract

Embryos develop in a concerted sequence of spatio-temporal arrangements of cells. In the preimplantation mouse embryo, the distribution of the cells in the inner cell mass evolves from a salt-and-pepper pattern to spatial segregation of two distinct cell types. The exact properties of the salt-and-pepper pattern have not been analysed so far. We investigate the spatio-temporal distribution of NANOG and GATA6 expressing cells in the ICM of the mouse blastocysts with quantitative three-dimensional single cell-based neighbourhood analyses. A combination of spatial statistics and agent-based modelling reveals that the cell fate distribution follows a local clustering pattern. Using ordinary differential equations modelling, we show that this pattern can be established by a distance-based signalling mechanism enabling cells to integrate information from the whole inner cell mass into their cell fate decision. Our work highlights the importance of longer-range signalling to ensure coordinated decisions in groups of cells to successfully build embryos.

## INTRODUCTION

During the development of multicellular organisms, coordinated cell behaviour is essential to shape the embryo. Cellular differentiation, proliferation and rearrangement is precisely regulated to obtain the correct positioning of the cells. This results in a variety of stable and reproducible patterns across the animal kingdom ^1–3^.

The mammalian preimplantation embryo exhibits distinct shape changes and patterns, some of which are conserved among different groups of Eutheria ^4^. After fertilisation, several cell divisions without cell growth yield a collection of smaller and smaller cells. Upon compaction, this loose cellular aggregate transforms into a compact, ball-like structure, and the embryo is referred to as a morula. At the same time, the outside cells of the morula develop an apical-basal polarity. The outside cells gradually form a fluid-tight seal and develop into an epithelium. This marks the first cell fate decision into the inner cell mass (ICM) which is enveloped by the outer trophectoderm (TE) that after implantation gives rise to the embryonic part of the placenta. Subsequently, inside the TE, next to the ICM, a fluid-filled cavity arises and the whole structure is known as blastocyst.

The cells in the ICM differentiate into epiblast (Epi) and primitive endoderm (PrE) precursors that after implantation give rise to the embryo and the yolk sac, respectively. The main markers used for the two lineages are NANOG for the Epi and GATA6 for the PrE. In early blastocysts, ICM cells co-express these markers. During differentiation in mid blastocysts, cells asynchronously downregulate NANOG or GATA6 to form PrE or Epi, respectively ^5^.

Besides marker expression, there is also cell positional changes ^6–8^. At the late blastocyst stage, this pattern is resolved by segregation of the two lineages such that the PrE forms a second epithelium at the surface of the ICM, separating the Epi cells from the cavity. The main pathway involved in the process is FGF/MAPK signalling, which reinforces PrE commitment: epiblast progenitors secrete FGF4, which binds to FGFR1 on Epi, and FGFR1 and FGFR2 on PrE biased cells ^9–15^. New analysis tools have been developed to study the neighbourhood structure in blastocysts such as IVEN ^16^. They reveal that an ICM cell has approximately 30% more neighbours than cells from the surrounding TE. Furthermore, the two cell types are clearly separated such that ICM cells have a majority of ICM neighbours and TE cells have mainly TE neighbours.

To investigate this process, we have previously used three-dimensional imaging of the mouse embryo with single cell resolution based on immunofluorescence staining, together with three-dimensional neighbourhood analyses ^17^. This approach produces cell positional information as well as cellular protein expression levels required to study the salt-and-pepper pattern. Our work allowed us to propose that expression levels of NANOG and GATA6 in the ICM during blastocyst development show a transition from local patterns in early blastocysts to a global pattern in the late blastocysts ^17^.

In this study, we quantitatively characterise the salt-and-pepper pattern in the ICM and identify a potential mechanism for the generation of the cell fate distribution. We classify the cells into binary groups of high or low expression levels of NANOG or GATA6. Comparison of the spatial cell distributions with *in silico* generated artificial realisations of the common notions of the salt-and-pepper pattern reveals that in early and mid blastocysts, the binary cell fates exhibit local clustering in the ICM. The key feature of this distribution is that the cellular composition of the direct neighbourhood of a cell corresponds to the cellular composition of the whole ICM.

Previous approaches have implemented nearest neighbour signalling under the consideration that the FGF/MAPK pathway is the key regulator of cell fate decisions in the ICM ^18–22^. We implemented two modelling approaches: nearest neighbour signalling as well as distance-based neighbour signalling and extended them to include cell division. Comparison of the spatial cell fate distributions between the simulations and the experimental data from our experiments (this study and ^17^ and other data sets ^5, 20^) reveals that cell signalling across the whole ICM best reproduces the spatial cell distribution observed experimentally.

## RESULTS

### Compiling available mouse preimplantation embryo data for neighbourhood analyses

We combined data from previous publications ^5, 17, 20^ with newly acquired embryo data (see M&M for details). This resulted in six data sets with 721 embryos of which 326 were early blastocysts (32-64 cells), 218 mid blastocysts (65-90 cells) and 177 late blastocysts (more than 90 cells). The analysis of the ICM population composition using NANOG (N) and GATA6 (G6) as fate markers shows the well-known trend for N-G6-, N+G6+, N-G6- and N-G6+ cell proportions (Fig. 1A, see also Sup. Fig. 1-6 for individually analysed data sets). In early blastocysts, most cells express both NANOG and GATA6. As development progresses, the proportion of N+G6+ cells decreases, while the proportion of N+G6-cells as well as N-G6+ cells increases. The N-G6-cell proportion is not so variable.

**Figure 1:**
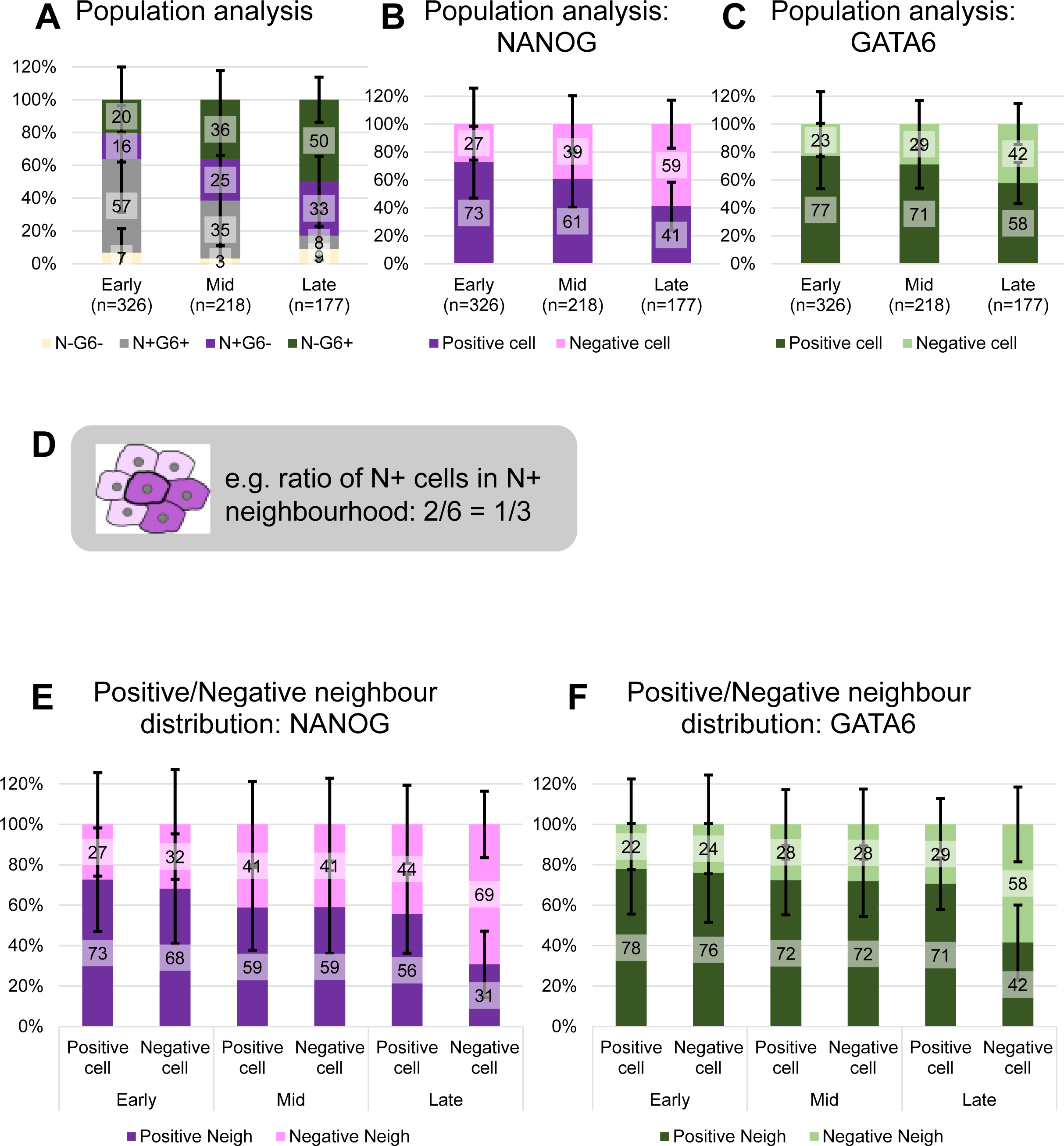
Average population and neighbourhood composition of all embryos in all data sets. **(A)** Results of ICM population analysis indicating the average percentage of N-G6-, N+G6+, N+G6- and N-G6+ cells in early (n=326), mid (n=218) and late (n=177) blastocysts. Error bars show the standard deviation, here and in subsequent graphs. **(B-C)** Results of ICM population analysis indicating the average percentage of N+ or N-cells (B) and G6+ or G6-cells (C) in early, mid and late blastocysts. **(D)** Neighbourhood analysis description scheme: the central cell in N+ and has two N+ neighbouring cells out of a total of 6 neighbours, hence 1/3 of its neighbourhood is composed of N+ cells. **(E-F)** Results of neighbour composition analyses for early, mid and late blastocysts indicating the average percentage of N+ or N-neighbours (E) and G6+ or G6-neighbours (F) of each ICM cell type. See Sup. Figs 1-6. for individually analysed data sets.

To investigate the nature of the salt-and-pepper pattern, it is essential that we classify the expression levels of NANOG and GATA6 independently (Fig. 1B-C). First, we introduced the categories NANOG+ and NANOG-cells (Fig. 1B, see M&M). The assignment of the ICM cells to these categories occurred independently of their respective GATA6 levels. Our results show that the proportion of NANOG+ cells in the ICM decreases from around 75% in early blastocysts to below 50% in late blastocysts. Categorising the cells only according to their GATA6 levels shows a decrease in the proportion of GATA6+ cells from 75% in early to 60% in late blastocysts (Fig. 1C).

Next, we analysed the composition of the local neighbourhood (Fig. 1D-F). The cell neighbourhood composition analysis gives information about each cell type and their neighbours. Hence, we considered the mean ratio per embryo of positive neighbours of positive cells (see illustration, Fig. 1D). Analogously, we included the mean ratio per embryo of negative neighbours of negative cells. For NANOG, we find that both NANOG+ and NANOG-cells have mostly NANOG+ neighbours in early and mid blastocysts. In late blastocysts, we observe a clustering such that NANOG-cells have mainly negative neighbours and NANOG+ cells have mainly positive neighbours (Fig. 1E). For GATA6, we see a similar behaviour (Fig. 1F). In early and mid blastocysts, all cells have mostly GATA6+ neighbours, while in late blastocysts, GATA6+ cells cluster with GATA6+ cells and GATA6-cells cluster with GATA6-cells.

In summary, we compiled several single-cell imaging data sets of mouse embryos to increase the reliability of the analyses results. We find that the binary cell fates NANOG+ and GATA6+ show a clustering in late embryos. Furthermore, the summary statistics are characterised by a large variability between individual embryos from the same stage that remains if we consider the data from each experimental study independently (Supp. Fig. 1-6).

### The local cell neighbourhood and global ICM population composition are strongly correlated

We have previously shown the importance of local cell neighbourhood during cell fate acquisition in mouse blastocysts ^17^. Others have highlighted the importance of the cell fate proportions in the ICM ^5, 20^. Hence, we wondered how the local cell neighbourhood is related to the ICM population composition. To investigate that, we checked if there is a relation between the neighbours’ composition and the overall proportion of positive and negative cells in each individual embryo (Fig. 2A-B, Supp. Fig. 7). To do this, we generated ICM composition scatter plots in which we represent the mean ratio of neighbours of a specific cell type per embryo versus the total proportion of the same cell type in the whole ICM in that embryo (i.e., mean proportion of positive neighbours of a positive cell versus total positive cells in the ICM). Indeed, we find that in early and mid blastocysts, for NANOG and GATA6, for positive as well as negative cells, the average proportion of the same cell type in the direct neighbourhood shows a strong correlation with the proportion of that cell type in the ICM. Furthermore, the point lies on the first bisector, indicating a proportionality with coefficient 1. Thus, if e.g. an embryo has 20% NANOG+ cells, the NANOG+ cells will have an average of 20% NANOG+ neighbours. For late embryos, the situation changes, as the correlation decreases. The scatterplot shows a cloud structure that is denser than for early and mid embryos. Hence, the proportions of positive and negative cells in the ICM as well as the proportions of neighbours are less variable. Furthermore, these neighbourhood proportions are larger than the proportion of that cell fate in the respective ICM. These results agree with the clustering of epiblast and primitive endoderm cells just before the embryo implants.

**Figure 2:**
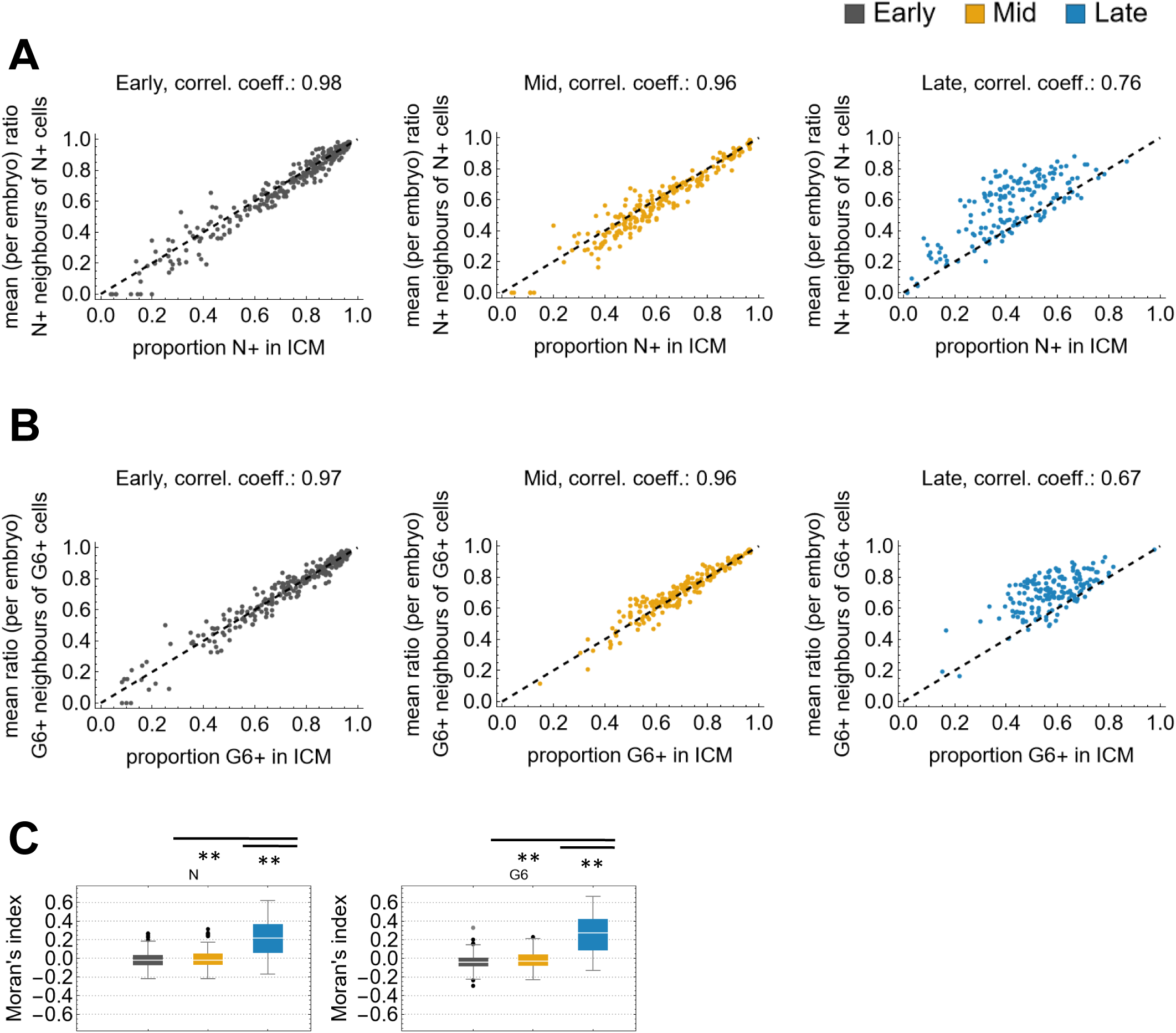
Single embryo neighbourhood quantification. **(A-B)** ICM composition scatter plot showing the mean ratio of N+ neighbours of N+ cells against the proportion of N+ in each ICM (A) and the mean ratio of G6+ neighbours of G6+ cells against the proportion of G6+ in each ICM (B) in early (left, grey), mid (middle, yellow) and late (right, blue) blastocysts. Each dot represents one embryo. See Sup. Fig. 7 for the plots of negative cells. Pearson’s correlation coefficient is indicated above each plot. **(C)** Box-Whisker-Charts of Moran’s indices for the patterns of NANOG (left) and GATA6 (right) positive and negative cells for all data sets for early (grey), mid (yellow) and late (blue) blastocysts. The stars indicate significant differences at a significance level of 0.01 obtained by a Welsh’s t-test with Bonferroni correction.

Since we are interested in the spatial distribution of the different cell types in relation with their neighbours, we decided to calculate Moran’s index (Fig. 2C, ^23^). Moran’s index is a coefficient that measures the spatial autocorrelation within a data set and has previously been applied in other biological contexts like the spatio-temporal patterning of brain activities or cancer ^24, 25^. In our context of cell fate decisions, it measures how similar a cell’s fate is to that of its neighbours. A value close to -1 means that the neighbouring cells are of opposite type (checkerboard or period two pattern). If cell fates form large clusters, Moran’s index will be positive, and it tends to 1 for a clear separation of two cell fates (sorted, see M&M). We got median Moran’s indices close to zero for NANOG and GATA6 in early and mid blastocysts: NANOG early (Ne): -0.01, NANOG mid (Nm): -0.001, GATA6 early (Ge): -0.05, GATA6 mid (Gm): -0.04 (Fig. 2C). The Moran’s index for late blastocysts is larger, consistent with the segregation of positive and negative cells.

In, summary, our results in late embryos show signs of tissue separation. For early and mid embryos, we find that the spatial cell fate distributions are stable during development. Interestingly, for each individual embryo, the local neighbourhood composition is strongly correlated and comparable to the population composition of the whole ICM. Different to previous assumptions ^18^, the Moran’s index does not indicate a checkerboard in early and mid embryos. Now, the question arises what kind of spatial cell fate pattern best describes these distributions.

### NANOG and GATA6 expressing cells show characteristics of local clustering in early and mid blastocysts

To improve our understanding of the cell fate distributions in the ICM, we employed our previously established rule-based simulations of artificial patterns (Fig. 3 and Sup. Fig.8-9, ^26^).

**Figure 3:**
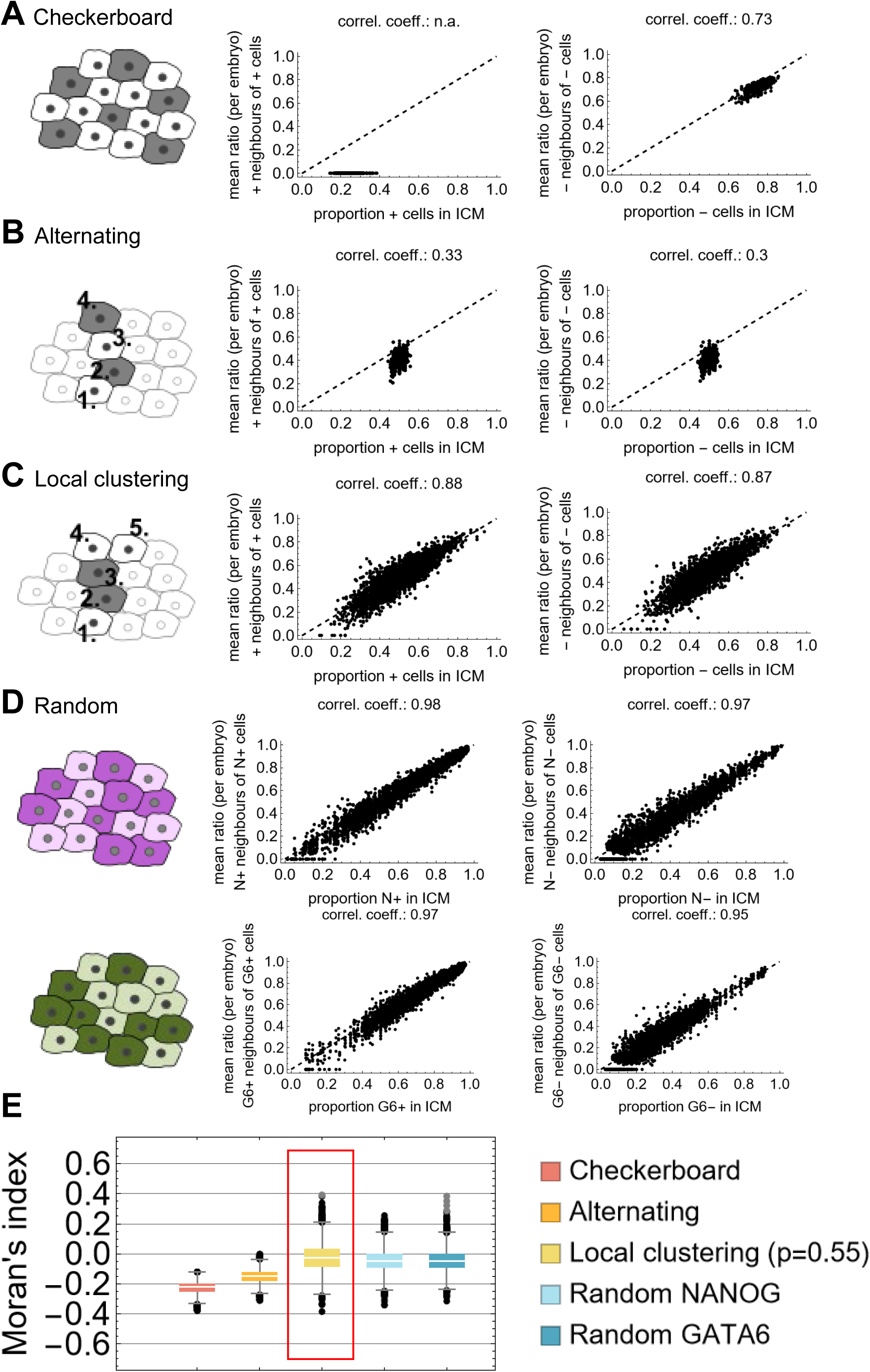
Single embryo neighbourhood rule-based models. **(A-D)** ICM composition scatter plot for the simulation results. The first column shows a two-dimensional scheme for the checkerboard (A), alternating (B), local clustering (C) and random (D) models. The second column shows the scatter plots showing the mean ratio of positive neighbours of positive cells against the proportion of positive in each model ICM. The third column shows the plots for negative cells. Each dot represents one simulation for one ICM. Pearson’s correlation coefficient is indicated above each plot. Please note that for the neighbourhood composition of positive cells in the checkerboard pattern, Pearson’s correlation coefficient is not defined. See Sup. Fig. 8 for the plots separated by stage. See Sup. Fig. 9 for the population analysis plots for the rule-based model patterns. **(E)** Box-Whisker-Charts of Moran indices for the rule-based model patterns checkerboard, alternating, local clustering and random. For values separated by stage, see Sup. Fig. 10. For the model results in the red box, we could not detect a significant difference to the experimental data for the four cell types N+, G6+, N-, G6- in early and mid blastocysts. This is based on a significance level of 0.01 and Welch’s t-test with Bonferroni correction.

We used the geometry of the embryo data, i.e. the cell numbers and coordinates, and the following rules (see M&M for full information about the models’ implementation):

- Checkerboard (Fig. 3A): positive cells have only negative neighbours.
- Alternating (Fig. 3B): cell fate is alternating between nearest neighbour cells, i.e. if a cell is positive, the cell that is closest with respect to Euclidean distance is negative and so on.
- Local cluster with probability *p* (Fig. 3C): one cell is randomly assigned a positive or negative state. The nearest neighbour, based on Euclidean distance, is assigned the same state as the previous cell with a given probability *p*. We used here *p*=0.55, based on a parameter scan (Sup. Fig. 10E and M&M).
- Random (Fig. 3D): In this case, we used the experimental cell numbers and coordinates as in the previous models, but also the proportion of experimentally observed fates (Fig. 1B and C). Then, these fates proportions were randomly distributed across the ICM of the given embryo.

The results of the simulated patterns of all models were plotted using the ICM composition scatter plots. The checkerboard rule generates a fate ratio of 25 ± 4% NANOG+ to 75 ± 4% NANOG- cells (mean ± standard deviation) (Fig. 3A, Supp. Fig. 8A and 9A). According to the pattern definition, the NANOG+ cell ratio in the neighbourhood of a NANOG+ cell is 0.

For the alternating model, we observe 50 ± 2% of positive and negative cells (Fig. 3B, Supp. Fig. 8B and 9B). In these two models, the results are independent of the embryo stage (Sup. Fig. 8A-B).

Local clustering with *p*=0.55 yields proportions of positive and negative cells in the ICM of 50 ± 11% of positive and negative cells (Fig. 3C, Supp. Fig. 8C and 9C). Looking at individual stages reveals a slight decrease in the spread of cell type proportions as development progresses (Sup. Fig. 8C). The standard deviation decreases from 12% in early, to 11 % in mid, and 9% in late embryos. In addition, the ratio of same cells in a cells direct neighbourhood agrees well with the ratio of the respective fate in the whole ICM as observed experimentally (Fig. 3C).

In the random model, the ratio of cells with the same fate in the direct neighbourhood of a cell shows a very strong correlation with the proportion of that fate in the whole tissue (Fig. 3D, Supp. Fig. 8D and 9D-E). For very low proportions of NANOG+ or NANOG-cells in the whole tissue, there are a few outliers with zero cells of the same fate in the neighbourhood: these are usually due to only one or two cells of that fate which are isolated from each other in the tissue. These results are again mostly independent of the embryo stage.

Comparing the ICM composition scatter plots, we observe that in the checkerboard and alternating models, the points lie mostly below the first bisector. However, in the local clustering and random models, these mostly lie on the first bisector, better representing the experimental data.

To compare the modelling results with the experimental data quantitatively, we calculated Moran’s index (Fig. 3E). The Moran’s indices for the rule-based simulations for checkerboard, alternating and random are all similar across stages (Sup. Fig. 10A-D) and significantly different from the experimental data. For the Moran’s indices for the local clustering model with *p*=0.55, we could not detect a significant difference from the experimental data.

In summary, we find that the experimental data is best reproduced by a local clustering model with a clustering probability of *p*=0.55. Hence, the question is what kind of signalling mechanism during development could result in such a distribution.

### Three-dimensional cell neighbourhood analyses challenge the nearest neighbour signalling model for cell fate decisions in the ICM

Previous mathematical models of primitive endoderm differentiation rely in essence on a bi- or tristable switch in a cell combined with an intercellular signal between direct neighbours ^18–22^. To test the neighbourhood relations resulting from these models, we implemented a model for two hypothetical transcription factors u and v that inhibit each other in a cell and are also subject to auto-activation (Fig. 4A) ^27^. In addition, we include an intercellular signal s that depends on the levels of u in the cell neighbourhood. In the given cell, the signal has an inhibitory effect on u and an activating effect on v.

**Figure 4:**
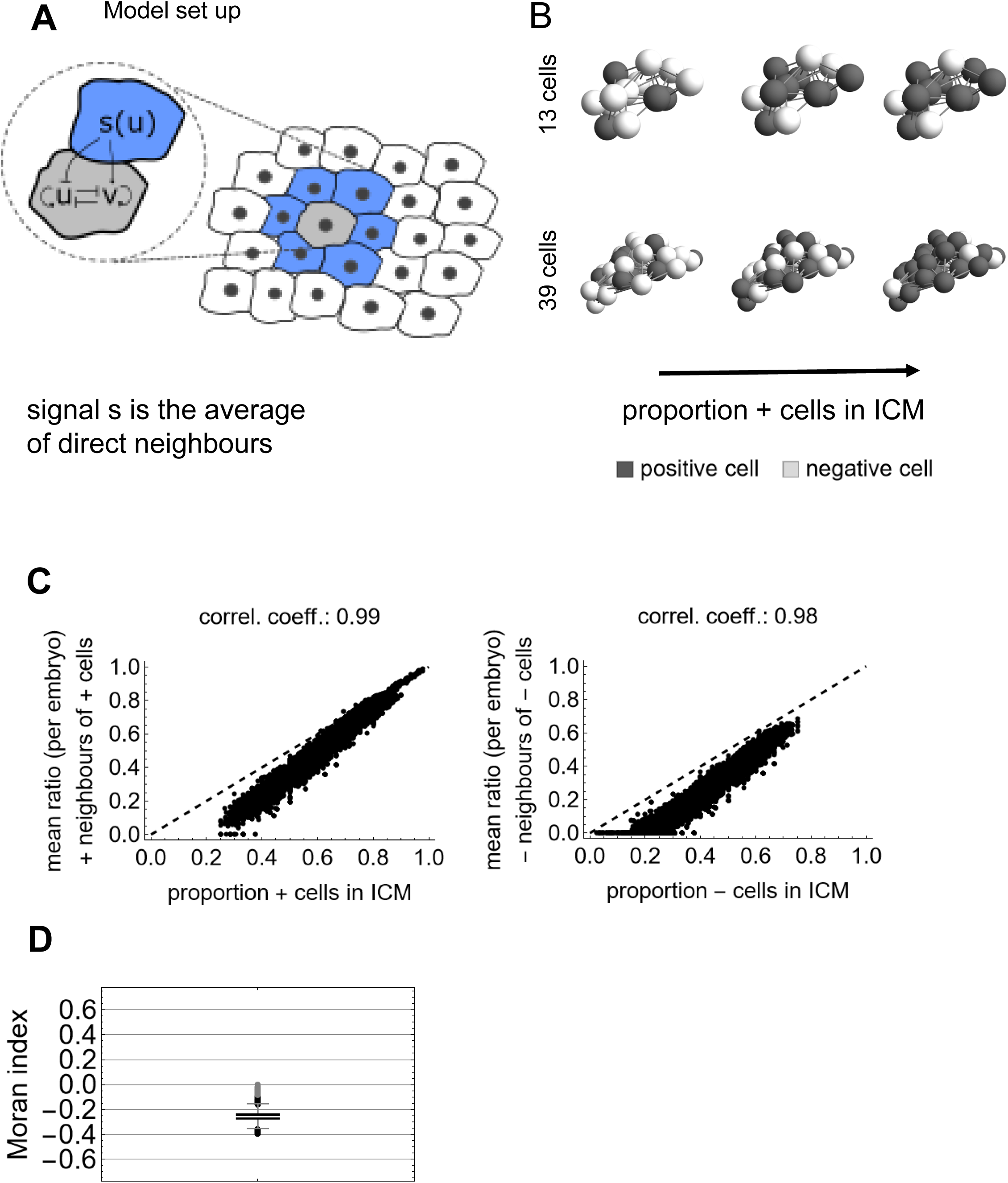
Nearest neighbour signalling: Model setup and simulation results. **(A)** Illustration of the gene regulatory network considered in the nearest neighbour signalling model. Inside the cell, u and v mutually inhibit each other. Additionally, v gets activated by an extracellular signal (s), whereas u is inhibited by the same. In this model, the extracellular signal to a given cell (grey cell in the scheme) is provided by its direct neighbours (blue cells). **(B)** Exemplary simulation results for three different cell fate ratios (30%, 60% and 80% positive cells) and two cell numbers. The top represents an ICM of 13 cells corresponding to an early blastocyst. The bottom one represents an ICM of 39 cells corresponding to a late embryo. **(C)** ICM composition scatter plot for the simulation results showing the mean ratio of positive neighbours of positive cells against the proportion of positive cells (left), and the mean ratio of negative neighbours of negative cells against the proportion of negative cells in each simulated ICM (right). Each dot represents one simulation for one ICM. Pearson’s correlation coefficient is indicated above each plot. **(D)** Box-Whisker-Chart of Moran’s indices for all simulations. See Sup. Fig 11 for results of the separation by stages.

As a first step, we assumed that the signal is given by the average levels of u of the nearest neighbouring cells (Fig 4A). To enable direct comparison between the simulations and the experimental data, we used ICM cell coordinates as the basis for our simulations. Furthermore, we set u-v+ as positive cell as and u+v- as negative cells. Example simulations show a trend towards a checkerboard-like pattern as in ^27^ although it is less clear because of the small number of cells and the three-dimensional arrangement (Fig. 4B). In the ICM composition scatter plots, the ratio of same fates in the direct neighbourhood lies clearly below the ratio of that fate in the whole tissue for both positive and negative cells (Fig. 4C, see Sup. Fig. 11A for results separated by stage). For high proportions of positive cells (>0.9), the ratio of positive cells in the neighbourhood of a positive cell is identical to the proportion of positive cells in the whole tissue. An interesting behaviour occurs for proportions of negative cells below 0.2. Here, the mean ratio of negative neighbours of negative cells is zero. Hence, negative cells do not have negative cells in their direct neighbourhood. The median Moran’s index is -0.26, again hinting at a checkerboard-like pattern (Fig. 4D). These results are independent of the stage, i.e., the number of cells (Sup. Fig. 11B).

These results show that nearest neighbour signalling cannot recapitulate the spatial cell fate distributions in mouse blastocysts. Hence, we extended the signalling mechanism to include cells beyond the nearest neighbours.

### A model in which ICM cells integrate signalling from beyond the nearest neighbours recapitulates the experimental spatial cell distribution

We assumed that cells beyond the nearest neighbours contribute to the signal for a given cell. We previously suggested a distance-based signalling mechanism and investigated its potential for spatial pattern formation (Fig. 5A, ^27^). A dispersion parameter *q*, with values between 0 and 1 regulates how strongly the intercellular signal propagates in the *in silico* tissue. For small *q*, cells interact mainly with their direct neighbours For *q*=0.1, e.g. 90% of the signalling reaches the first neighbours, while cells further away receive only 10% of the signal. In comparison, for *q*=0.5 the signal is halved every time it reaches cells one graph distance further away. Different values for *q* result in different patterns for the two cell types. On artificial tissues with 200 cells, small *q* provides a checkerboard-like pattern. Increasing *q* yields increasing cell fate clusters and for *q*=1 an engulfing pattern arises.

**Figure 5:**
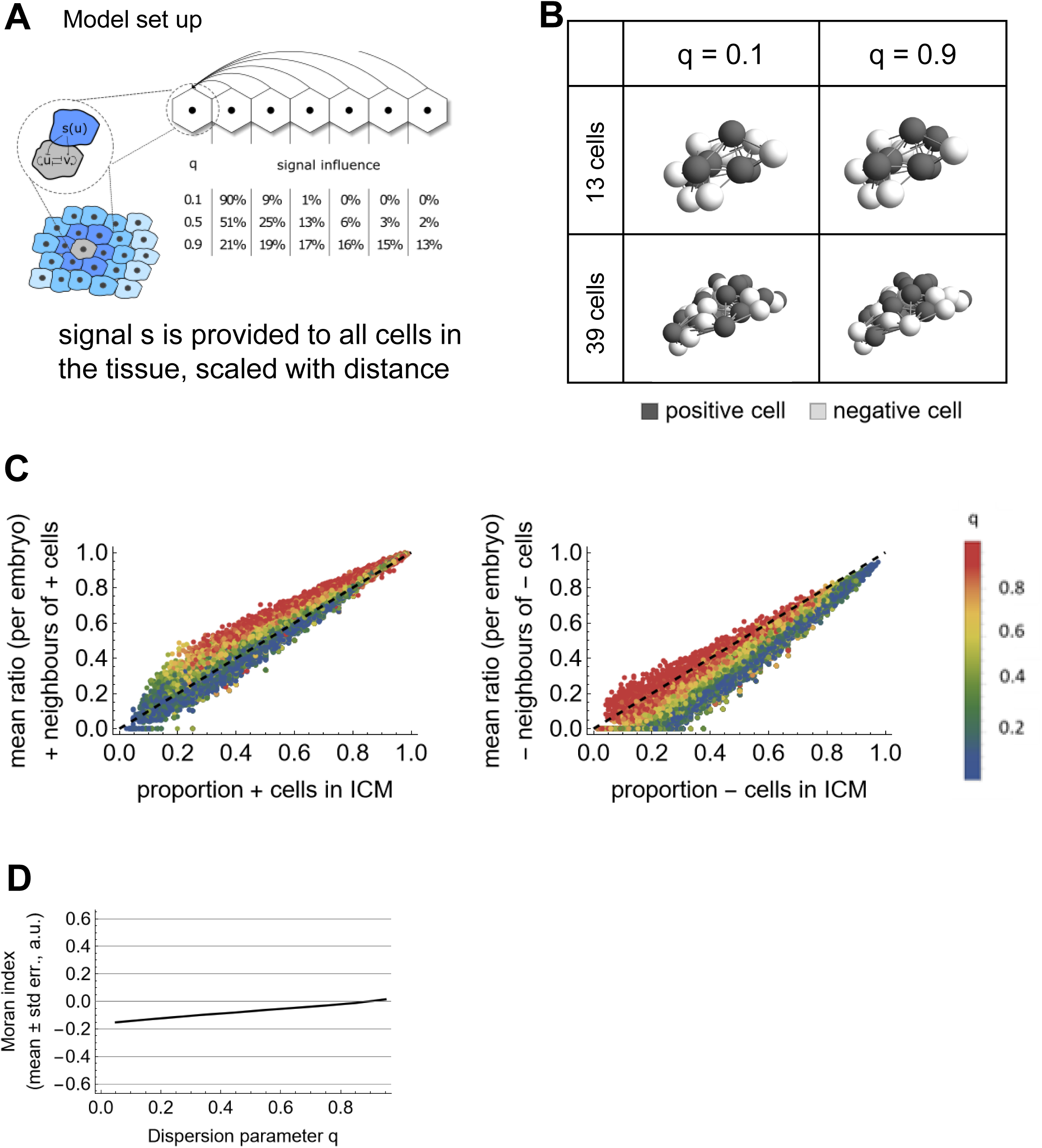
Distance-based neighbour signalling: Model setup and simulation results. **(A)** Illustration of the gene regulatory network considered in the distance-based signalling model. Inside the cell, u and v mutually inhibit each other. Additionally, v gets activated by an extracellular signal, whereas u is inhibited by the same. In this model, the extracellular signal is the sum of all cell-cell communication between one cell and any other cell in the system. The intensity of signalling contribution depends on the dispersion parameter q and the distance d_ij_ between the signal-receiving cell i and the signal-sending cell j. **(B)** Exemplary simulation results for two different values of the dispersion parameter q and two cell numbers (13 - an example for an early ICM and 39 – an example for a late ICM). The proportion of cells was set to 50% positive cells. **(C)** ICM composition scatter plot for the simulation results showing the mean ratio of positive neighbours of positive cells against the proportion of positive cells for each simulation for an ICM (left), and the mean ratio of negative neighbours of negative cells against the proportion of negative cells for each simulation for an ICM (right). The colour code indicates the *q* values. Each dot represents one simulation for one ICM. **(D)** Moran’s index against dispersion parameter q. The plot consists of a solid line for the average and a shaded area for the standard error of the mean (SEM). Please note that the SEM is very small and therefore hardly visible.

**Figure 6:**
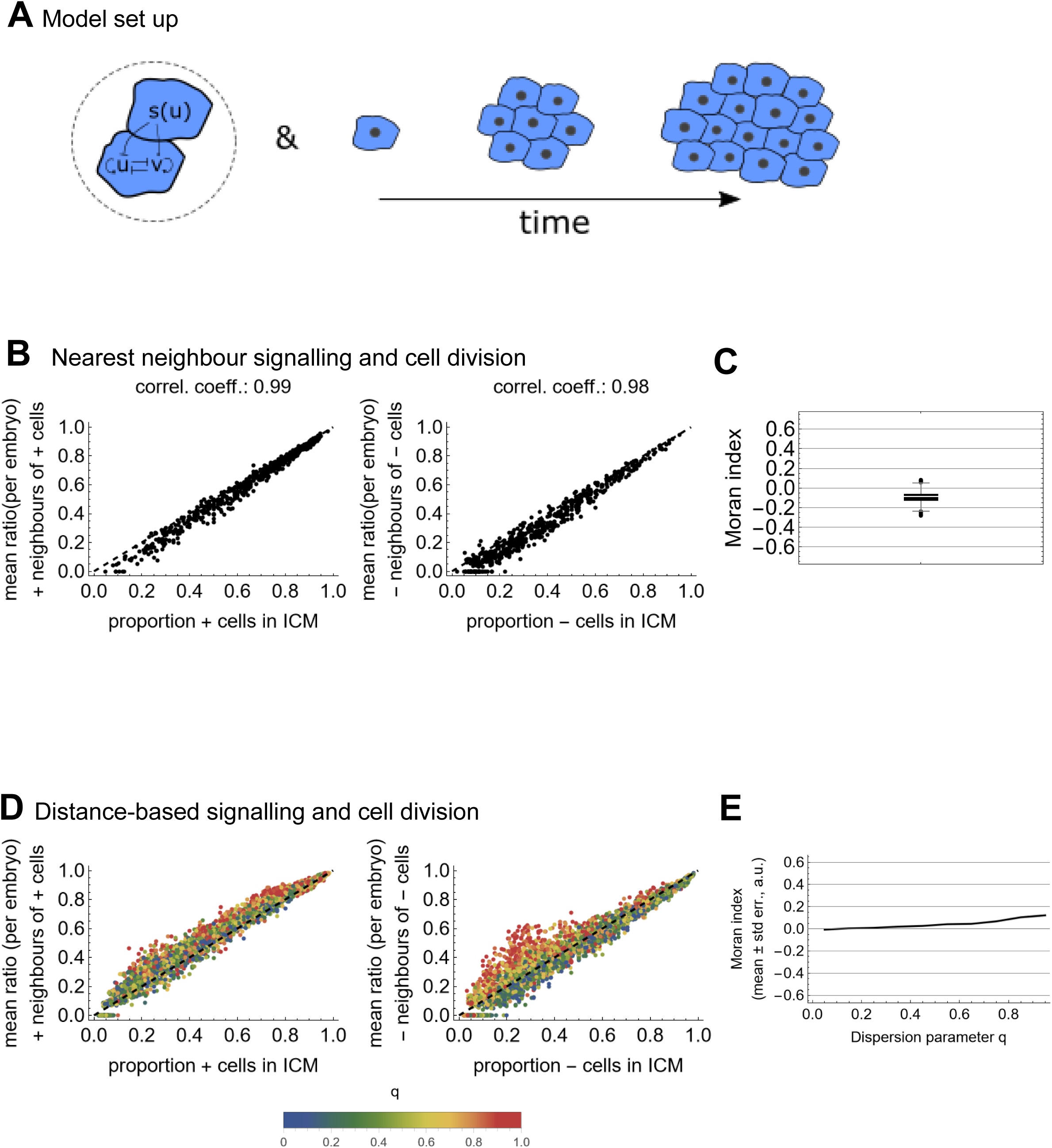
Nearest neighbour and distance-based neighbour signalling with cell division: Model setup and simulation results. **(A)** llustration of the combination of a neighbour signalling model with cell division. **(B)** ICM composition scatter plot for the simulation results showing the mean ratio of positive neighbours of positive cells against the proportion of positive cells in each simulated ICM (left), and the mean ratio of negative neighbours of negative cells against the proportion of negative cells in each simulated ICM (right). Each dot represents one simulated ICM. Pearson’s correlation coefficient is indicated above each plot. **(C)** Box-Whisker-Chart of Moran’s indices for all simulations. For separation by stages, see Sup. Fig 13. **(D)** ICM composition scatter plot for the simulation results showing the mean ratio of positive neighbours of positive cells against the proportion of positive cells in each simulated ICM (left), and the mean ratio of negative neighbours of negative cells against the proportion of negative cells in each simulated ICM (right). The colour coding indicates the *q* values. Each dot represents one simulation for one ICM. **(E)** Moran’s index against dispersion parameter q. The plot consists of a solid line for the average and a shaded area for the standard error of the mean (SEM). Please note that the SEM is very small and therefore hardly visible. For separation by stages, see Sup. Fig 13.

**Figure 7:**
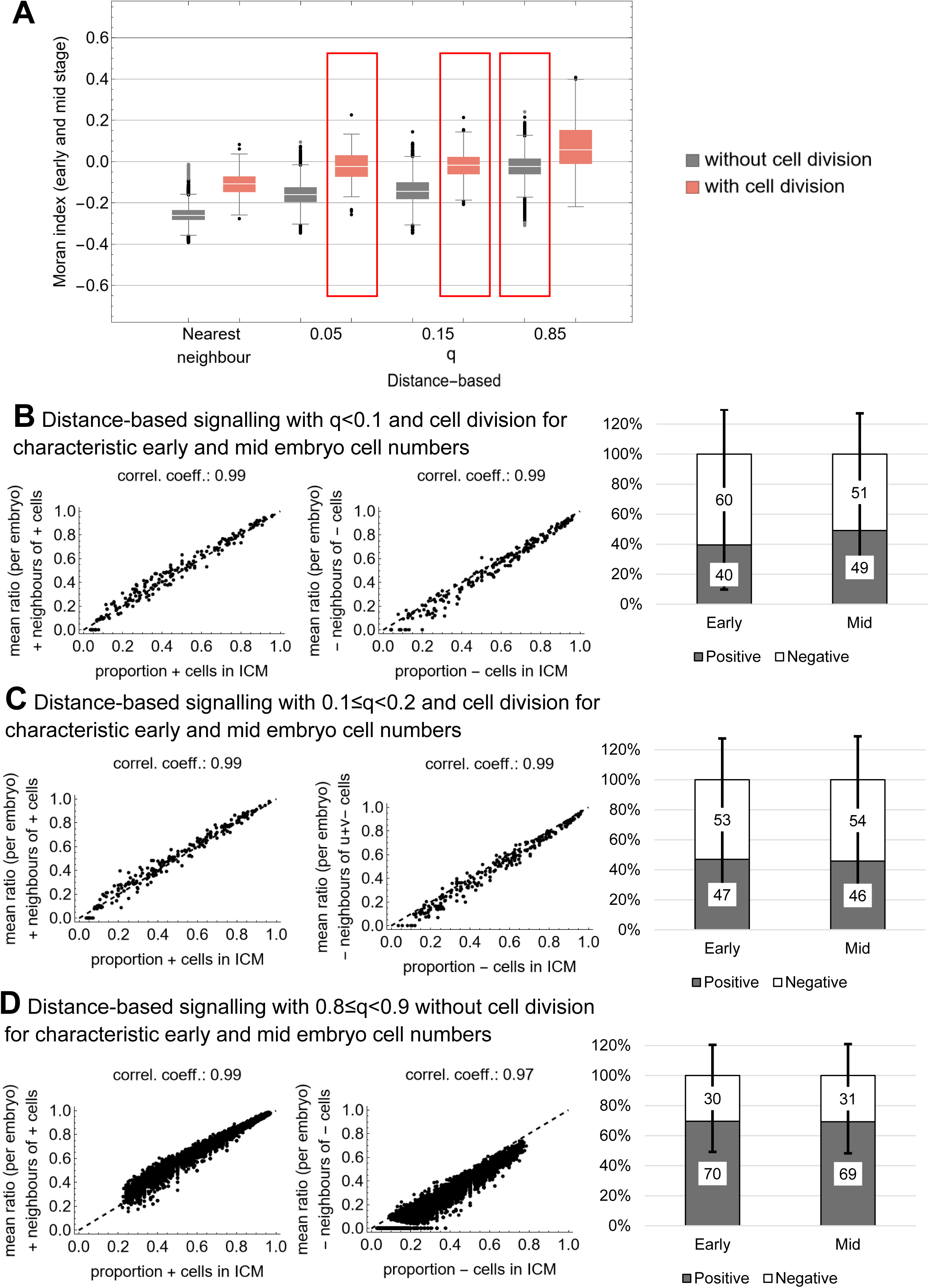
Best fitting signalling models. **(A)** Box-Whisker-Chart of Moran’s indices for indicated simulations. Box-Whisker-Charts of Moran’s indices for nearest neighbour and distance-based neighbour signalling without (grey) and with (light red) cell division. The distance-based neighbour signalling models were collected in bins of width 0.1 based on the respective q value. The bin centres are used as labels. For the model results in the red box, we could not detect a significant difference to the experimental data for the four cell types N+, G6+, N-, G6- in early and mid blastocysts. **(B-C)** ICM composition scatter plots for the simulation results for the distance-based signalling model with *q*<0.1 (B) and 0.1≤*q*<0.2 (C) and cell division. Scatter plots showing the mean ratio of positive neighbours of positive cells against the proportion of positive cells for each simulation for an ICM (left), and the mean ratio of negative neighbours of negative cells against the proportion of negative cells for each simulation for an ICM (centre). Each dot represents one simulation for one ICM of early and mid embryos. (Right) ICM population analysis indicating the average percentage of positive and negative cells in the simulations for early, mid and late ICMs. **(D)** ICM composition scatter plot for the simulation results for the distance-based signalling model with 0.8≤*q*<0.9 and no cell division. Scatter plots showing the mean ratio of positive neighbours of positive cells against the proportion of positive cells for each simulation for an ICM (left), and the mean ratio of negative neighbours of negative cells against the proportion of negative cells for each simulation for an ICM (centre). Each dot represents one simulation for one ICM of early and mid embryos. (Right) ICM population analysis indicating the average percentage of positive and negative cells in the simulations for early, mid and late ICMs.

We performed simulations of this model on tissues that are determined by the coordinates of *in vivo* ICM cells. Example simulations on two ICMs with different cell numbers show an increase in clustering with increasing *q* although it is less obvious than for the larger simulated tissues (Fig. 5B, ^27^). The ICM composition scatter plots emphasize the changes in patterning with increasing *q* (Fig. 5C see Sup. Fig. 12A for results separated by stage). For positive cells and low *q*, the average neighbourhood composition corresponds to the overall ratio of positive cells in the individual ICMs. As *q* increases, the curve moves above this line (Fig. 5C, left). For negative cells, the results lie below the first intercept for most *q*. We believe the main reason for this is that our mechanism has a radial bias such that negative cells tend to accumulate at the boundary of the tissue (Fig 5C, right). Moran’s index supports these findings (Fig. 5D, Sup. Fig. 12B). For low *q*, it is negative and increases with increasing *q*. For *q* above 0.8, Moran’s index is around zero. This holds true if we separate the simulated data by the stages of the underlying embryo geometries (Supp Fig 12B).

In summary, the observed spatial cell fate distribution is compatible with a model in which ICM cells integrate signalling across a longer range. In early and mid ICMs, this range can be up to 5 cells, in late embryos up to 8 cells (Sup. Fig. 12C).

### Including cell division improves the fit of the spatial cell fate distributions for shorter-range signalling

We have previously proposed cell division as a mechanism that generates clusters in cell fate patterns ^28^. Therefore, we combined the two intercellular signalling mechanisms with cell division and analysed the resulting spatial patterns (Fig. 6A).

The ICM composition scatter plots for the model with nearest neighbour signalling that includes cell division show that the local neighbourhood distribution gets closer to the proportion of the respective cells in the tissue and the median Moran’s index increases to -

0.09 (Fig. 6B-C and Sup. Fig. 13A-B). Separating the results by the stage of the embryo geometry shows a slight increase of the Moran’s index from –0.1 for early and mid to 0.06 for late blastocysts (Supp. Fig. 13B). The results indicate that a model that includes nearest neighbour signalling with cell division produces a cell fate pattern which better resembles the experimental data than the model without division. However, the population composition does not correspond to the observed one (Supp. Fig. 13C).

The inclusion of cell division in the model with distance-based signalling shifted the results such that a distribution on the first bisector with Moran’s index zero is now obtained for smaller values of *q* (Fig. 6D-E and Sup. Fig. 13D-E). Separating the data by the stage of the ICM geometry also shows a stage dependence (Supp. Fig. 13D-E). Only early and mid ICM geometries can exhibit a fate distribution on the first bisector with Moran’s index zero. For late ICM geometries, the neighbourhood ratios are larger than the respective cell fate proportions in the ICM and the Moran’s index is larger than zero for most *q*.

In summary, including cell division in the nearest neighbour signalling model reduces the discrepancy between simulation results and experimental data. However, they still do not match the observed spatial cell fate patterns sufficiently well. Adding cell division to the distance-based model reduces the required signalling distance during cell fate decision to recapitulate the observed spatial patterns.

### The spatial patterns of NANOG and GATA6 expressing cells in early and mid blastocysts are consistent with a mechanism that integrates intercellular signalling from the whole ICM without cell division

To determine the best fitting signalling model, we compared the Moran’s indices of the different models for early and mid blastocysts (Fig. 7). To this end, we binned the Moran’s indices for the distance-based models according to their *q* values in 10 bins of width 0.1 with bin centres at 0.05, 0.15, etc. We found three models for which we could not detect a statistically significant difference to the experimental data for NANOG and GATA6 patterns in early and mid embryos: distance-based signalling with *q*=0.05 and *q*=0.15 and cell division as well as distance-based signalling with *q*=0.85 and no cell division. For the ICM composition scatter plots, we see that both models show a strong correlation between the local neighbourhood composition and the ICM composition (Fig. 5B-C and Sup. Fig. 14). Furthermore, the points lie on the first bisector.

Looking at the cell fate ratios however provides a clear distinction (Fig. 7B left and 7C left). The model with cell division generates between 60-50% positive and 40-50% negative cells and no neighbour clustering effect of any cell type. However, the model without cell division results in 70% positive and 30% negative cells and clustering of positive cells. The latter is closer to the observed experimental data (Fig 1).

Overall, we conclude that a signalling mechanism that integrates the signals from all ICM cells with no cell division can yield the local cell fate clusters in mouse ICMs and additionally reproduce the population proportions observed in the ICM.

## DISCUSSION

It has been proposed that during cell fate acquisition in early mouse preimplantation embryos, Epi and PrE progenitors are located following a salt-and-pepper pattern. Traditionally, this pattern has been assumed to be random ^6–8^ or checkerboard-like ^18^ without investigating its true nature. Rather, cell fate acquisition has been mainly studied focusing on cell fate ratios with FGF/MAPK signalling as the main pathway involved in the process ^19^. However, other signalling pathways are also involved, modulating the cell fate acquisition process ^29^. Very few studies have considered the spatial cell fate distribution in their signalling models for the embryo ^18, 19^, and always without quantitative experimental data for comparison. In this study, we focused on understanding the salt-and-pepper pattern in early mouse preimplantation embryo. We integrated cell fate ratios, three-dimensional cell fate distribution and signalling. For this, we used single cell immunofluorescence imaging data of early, mid and late blastocysts, neighbourhood analyses, and rule-based modelling as well as simulations based on ordinary differential equations (ODEs) for a toggle switch coupled to intercellular signalling. We show that the salt- and-pepper pattern in mouse blastocyst ICMs is best represented by a local clustering pattern resulting from a signalling mechanism with a large dispersion range and no cell division. This result agrees with previous findings in three- and two-dimensional *in vitro* cell cultures by us and others us, respectively ^26, 30^. Our previous work with ICM organoids has revealed that the salt-and-pepper pattern of the *in vitro* system fits a local clustering pattern ^26^ and that predicting the fate of a given cell based on the neighbouring fates using neural networks only shows a 70% accuracy ^31^. Raina et al. observed a clustering effect in 2D in vitro cell cultures, which they could fit well by a model that considers nearest and second nearest neighbours for the signalling mechanism ^30^.

Previous ODEs models for cell fate decision in early mouse embryos have incorporated intracellular inhibition and intercellular signalling only between nearest neighbours in combination with cell division ^18, 20^. Tosenberger et al., analysed the three-dimensional cell fate distribution of the simulations, but have not presented experimental data on this part ^18^. Saiz et al., focused on cell fate ratios in late embryos. Cell fate ratios or the spatial distribution during differentiation were not considered ^20^. We implemented an analogous nearest neighbour signalling model with cell division and found that the simulation results for the spatial cell distribution were not comparable to the experimental data. Interestingly, our results further suggest that in early and mid embryos, not the absolute cell fate ratio, but rather the relation between the cell fate ratios in the neighbourhood and in the whole ICM are conserved.

In our study, the model showing the best fit with the large experimental data integrates signalling from nearest neighbours as well as from cells further away in the tissue, which have a considerable influence. Our analyses show that in early and mid ICMs, this can range to up to the fifth neighbour.

The main difference between nearest neighbour signalling and distance-based signalling is how far the signal travels. The main signal in this context is FGF4 ^11^ and it is usually assumed that it freely diffuses within the tissue. On a timescale of several days, FGF4 has been shown to diffuse over many millimetres in cell culture dishes and that embryonic stem cells use it for communication across these large scales ^32^. For a timescale of 12 h, FGF4 signalling reaches mainly first and second neighbours in embryonic stem cell cultures ^30^. FGF signalling range can be modulated by a number of mechanisms: FGF diffusion via heparan-sulphate proteoglycans at the extracellular matrix, FGFR trafficking and *fgf4* mRNA decay ^33^. All these cellular mechanisms have yet to be investigated in the developing mouse preimplantation embryo. Moreover, FGFs might not even diffuse at all, as has been shown for other morphogens like Wingless ^34^. In this context, it might also be possible that FGF signalling range is increased by cell cytonemes present in the ICM cells ^35^ or microvesicles ^36^. Furthermore, not only FGF4 is expressed in the early mouse embryo, other FGFs like FGF10, FGF3, and FGF5 are also expressed complicating the FGF/MAPK signalling scenario ^37^. Another cellular process that might have an effect in the cell neighbourhood and signal dispersion is the presence of cytoplasmic bridges connecting sister cells during embryonic development hence coordinating cell fate decisions ^38^. These bridges have been observed in mouse morulae ^39, 40^, as well as in Epi cells from postimplantation embryos ^41, 42^.

Surprisingly, the best fitting model in our study does not incorporate cell division. Cell division is currently implemented symmetrically since previous results for modelling the neighbourhood composition of ICM organoids have shown that symmetric division fits the experimental distributions best ^28^. In that regard, mouse embryonic stem cells expressing a *Nanog*-GPF reporter divide symmetrically during differentiation ^43^. However, an asymmetric distribution of the two transcription factors to the daughter cells could be plausible and might account for the differences in population distribution. Furthermore, a factor that we have neglected so far, is apoptosis. On average, 3.2±2.81 dead cells occur in the mouse ICM ^44^. Apoptosis mostly starts occurring in mid blastocysts and the rates steadily increase with increasing cell number ^6^. Hence, apoptosis could only account for the discrepancy for mid blastocysts. Furthermore, our model would predict that apoptosis of one cell fate dominates over the other. Our model currently predicts equal amounts of positive and negative cells. The experimental data shows a distribution of 30% negative and 70% positive cells both for NANOG and GATA6. To match this, an increase in cell death in negative cells would be required. Overall, a signalling model in which differential apoptosis and asymmetric cell division is implemented would be more biologically accurate. Finally, our model and local clustering pattern during cell fate acquisition in mouse preimplantation embryo is only applicable to early and mid blastocysts and does not fit late ones. This is likely due not only to enhanced cell division asymmetries or apoptosis, but also to cell migration of PrE cells to their final position facing the cavity before the embryo implants. Another factor influencing cell fate distribution in late blastocysts might be the FGF/MAPK signalling maturation, with changes in the expression levels of its components and activity ^12, 37^.

The notion of a salt-and-pepper pattern has also come up in other developmental questions like the cell fate segregation in pancreatic or neural development through Notch signalling ^45, 46^, the segregation of the definitive endoderm ^47^, mammalian visual cortex organization ^48^ or the cilia in the notochord ^49^. Focusing on the spatial statistics in the data analysis and the model fitting might also help to reveal the nature of the spatial pattern and improve the understanding of the underlying mechanisms in those contexts. In summary, using spatial statistics on single cell embryo data reveals that understanding cell fate decisions in mouse blastocysts must accommodate not only a cells neighbourhood, but also a larger proportion of the tissue. This ensures that cells make coordinated decisions to successfully make an embryo.

### Limitations of the study

The experimental data used in this study come from immunostained fixed embryos. Ideally the experimental data should have been obtained from time-lapse imaging data from double reporter mouse lines reporting NANOG and GATA6 protein levels. To our knowledge such mouse line is not currently available.

## Supporting information

Supplemental details

## Acknowledgements

Work at SMD lab was funded by the ACIISI (CEI2019-02), Programa de Ayudas a la Investigación de la ULPGC, and ACIISI co-funded by FEDER Funds (ProID2020010013). JLG is supported by the ULPGC predoctoral program. SMD was supported by the “Viera y Clavijo” Program from the Agencia Canaria de Investigación, Innovación y Sociedad de la Información (ACIISI) and the ULPGC. Work at the SF lab was supported through funding by the Deutsche Forschungsgemeinschaft (DFG, German Research Foundation) project number 470129398 and start-up funding by the University of Wuerzburg.

## Authors contributions

SCF developed the algorithms for data analysis, performed data analysis, supervised the work and wrote the manuscript; SS developed the model for intercellular signalling, performed the simulations and critically revised the manuscript; JLG did experimental work, mice maintenance, image analysis and critically revised the manuscript; SMD did experimental work, image analysis, data analysis, wrote the manuscript and supervised the work.

---

**Supplemental Figure 1:**
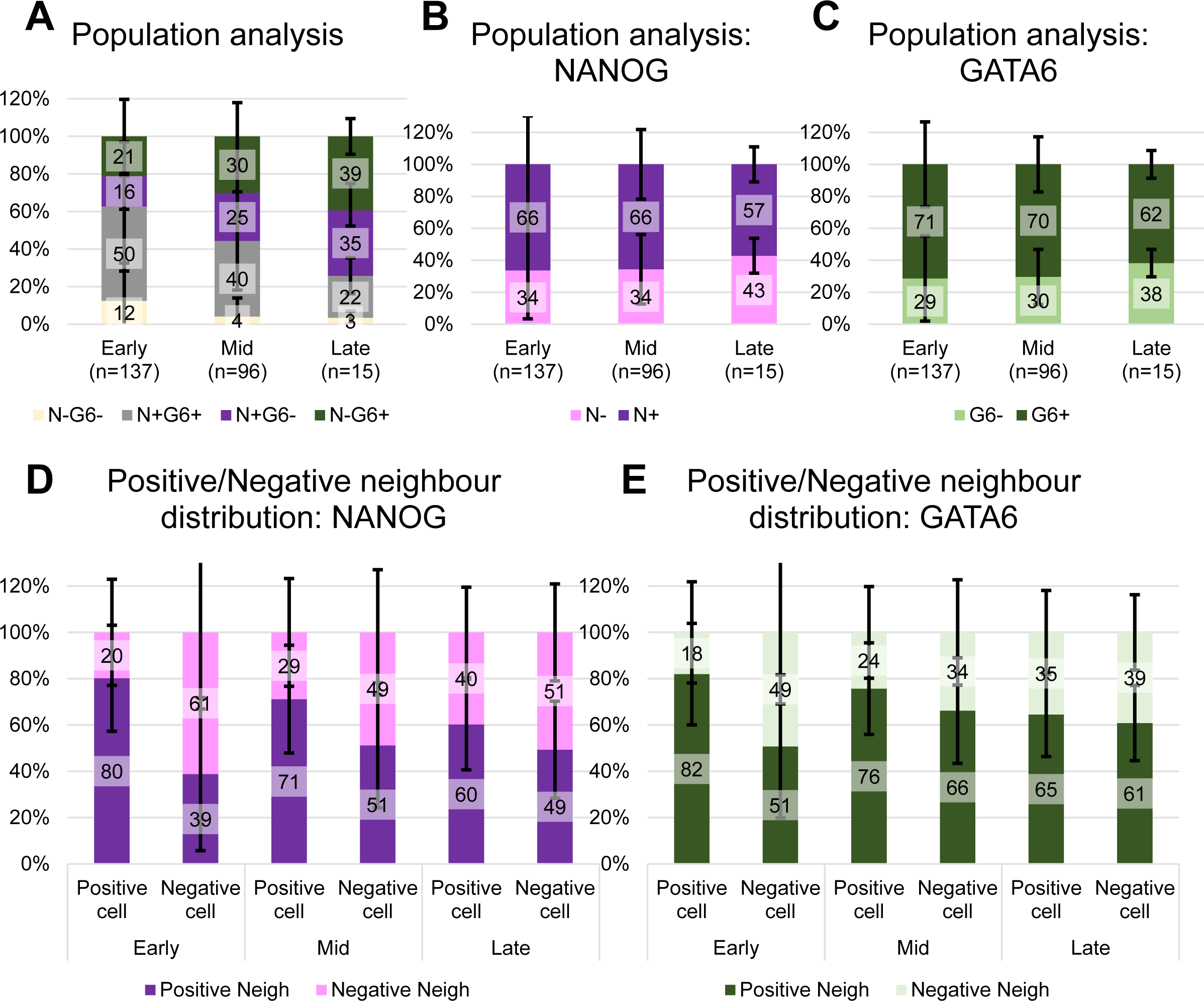
Average population and neighbourhood composition of all embryos in data set I. **(A)** Results of ICM population analysis indicating the average percentage of N-G6-, N+G6+, N+G6- and N-G6+ cells in early (n=137), mid (n=96) and late (n=15) blastocysts. Error bars show the standard deviation, here and in subsequent graphs. **(B-C)** Results of ICM population analysis indicating the average percentage of N+ or N-cells (B) and G6+ or G6- cells (C) in early, mid and late blastocysts. **(D-E)** Results of neighbour composition analyses for early, mid and late blastocysts indicating the average percentage of N+ or N-neighbours (D) and G6+ or G6-neighbours(E) of each ICM cell type.

**Supplemental Figure 2:**
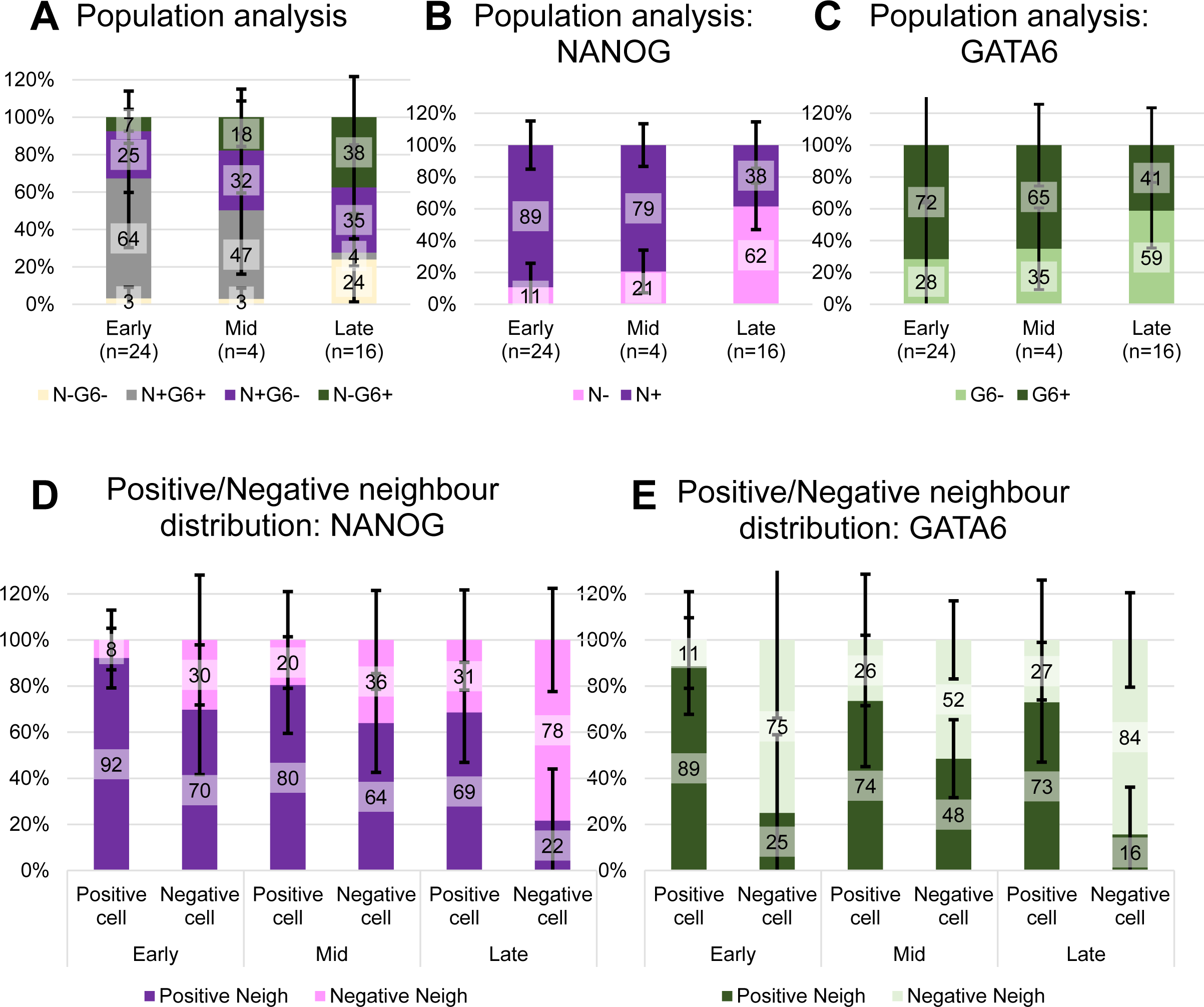
Average population and neighbourhood composition of all embryos in data set II^17^. **(A)** Results of ICM population analysis indicating the average percentage of N-G6-, N+G6+, N+G6- and N-G6+ cells in early (n=24), mid (n=4) and late (n=16) blastocysts. Error bars show the standard deviation, here and in subsequent graphs. **(B-C)** Results of ICM population analysis indicating the average percentage of N+ or N-cells (B) and G6+ or G6-cells (C) in early, mid and late blastocysts. **(D-E)** Results of neighbour composition analyses for early, mid and late blastocysts indicating the average percentage of N+ or N-neighbours (D) and G6+ or G6-neighbours(E) of each ICM cell type.

**Supplemental Figure 3:**
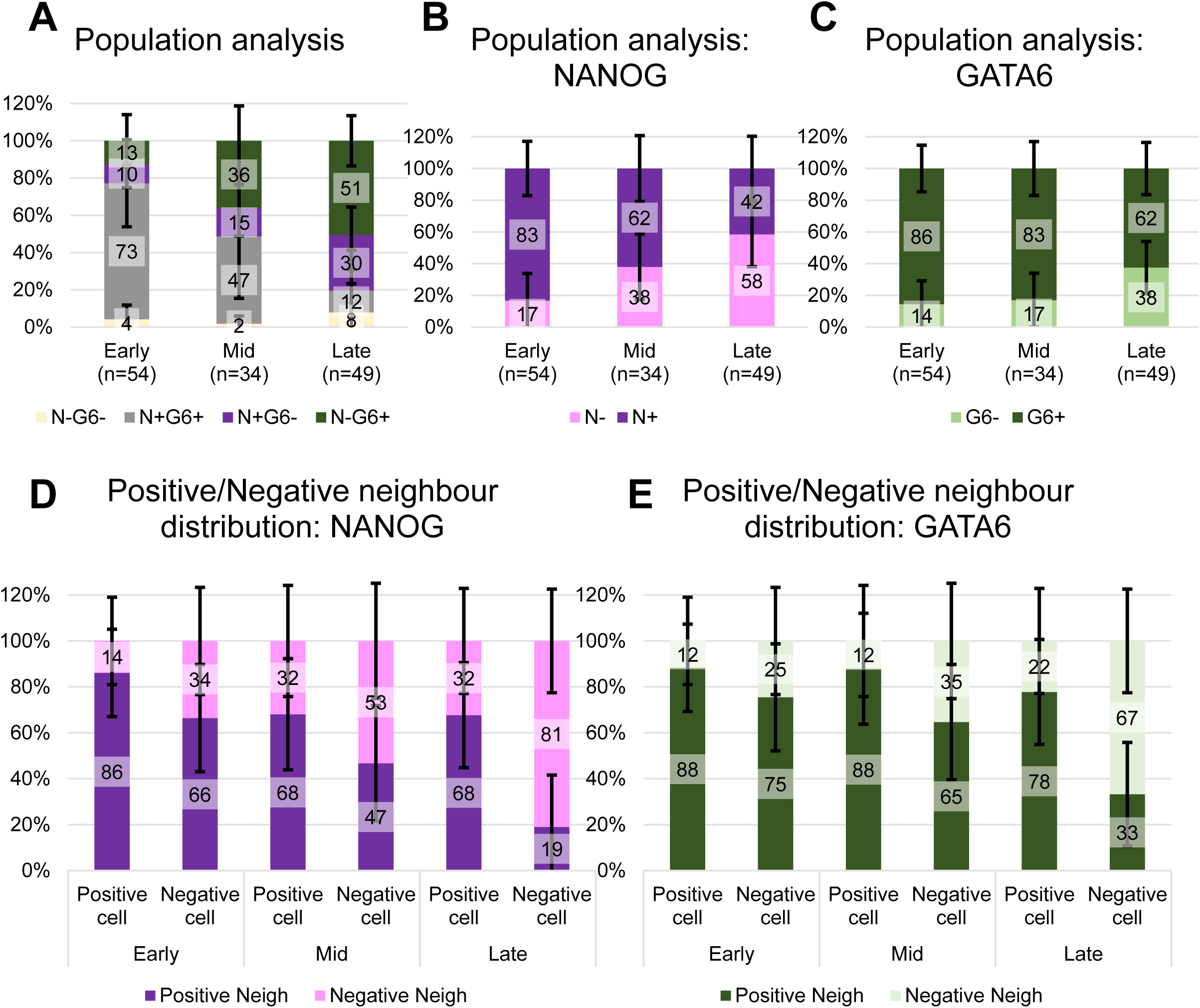
Average population and neighbourhood composition of all embryos in data set III^5^. **(A)** Results of ICM population analysis indicating the average percentage of N-G6-, N+G6+, N+G6- and N-G6+ cells in early (n=54), mid (n=34) and late (n=49) blastocysts. Error bars show the standard deviation, here and in subsequent graphs. **(B-C)** Results of ICM population analysis indicating the average percentage of N+ or N-cells (B) and G6+ or G6- cells (C) in early, mid and late blastocysts. **(D-E)** Results of neighbour composition analyses for early, mid and late blastocysts indicating the average percentage of N+ or N-neighbours (D) and G6+ or G6-neighbours(E) of each ICM cell type.

**Supplemental Figure 4:**
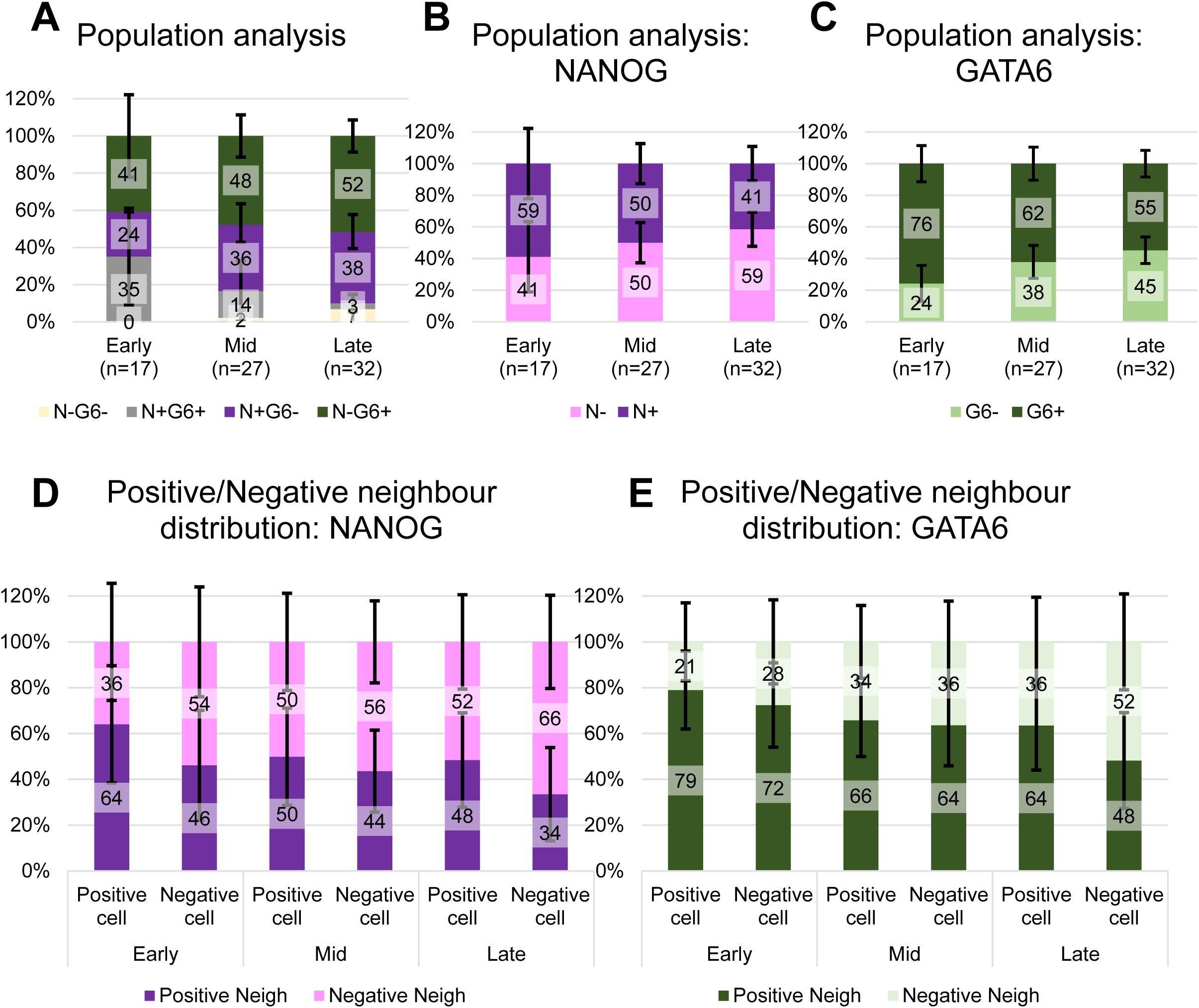
Average population and neighbourhood composition of all embryos in data set IV ^20^. **(A)** Results of ICM population analysis indicating the average percentage of N-G6-, N+G6+, N+G6- and N-G6+ cells in early (n=17), mid (n=27) and late (n=32) blastocysts. Error bars show the standard deviation, here and in subsequent graphs. **(B-C)** Results of ICM population analysis indicating the average percentage of N+ or N-cells (B) and G6+ or G6-cells (C) in early, mid and late blastocysts. **(D-E)** Results of neighbour composition analyses for early, mid and late blastocysts indicating the average percentage of N+ or N-neighbours (D) and G6+ or G6-neighbours(E) of each ICM cell type.

**Supplemental Figure 5:**
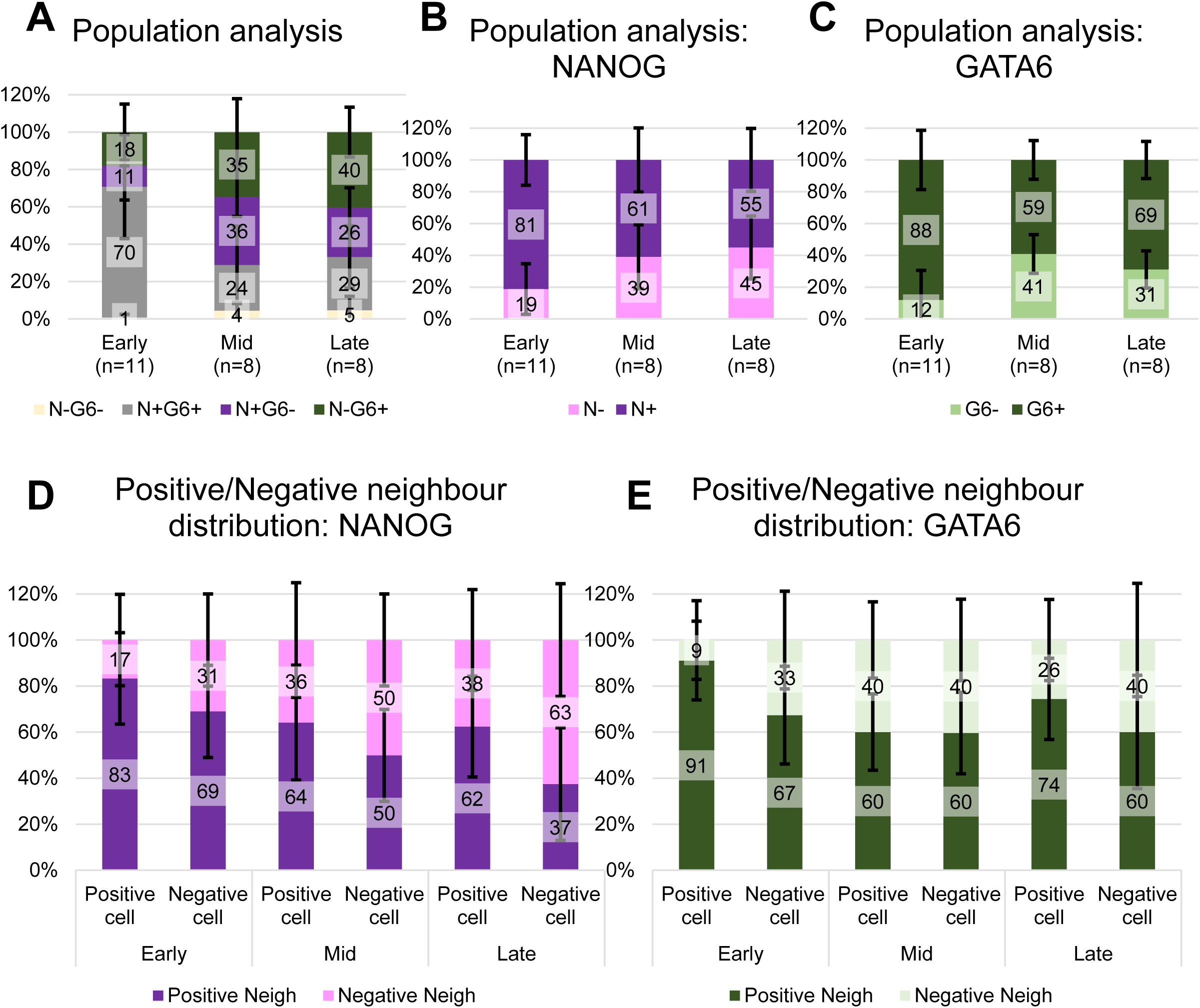
Average population and neighbourhood composition of all embryos in data set V^20^. **(A)** Results of ICM population analysis indicating the average percentage of N-G6-, N+G6+, N+G6- and N-G6+ cells in early (n=11), mid (n=8) and late (n=8) blastocysts. Error bars show the standard deviation, here and in subsequent graphs. **(B-C)** Results of ICM population analysis indicating the average percentage of N+ or N-cells (B) and G6+ or G6-cells (C) in early, mid and late blastocysts. **(D-E)** Results of neighbour composition analyses for early, mid and late blastocysts indicating the average percentage of N+ or N-neighbours (D) and G6+ or G6-neighbours(E) of each ICM cell type.

**Supplemental Figure 6:**
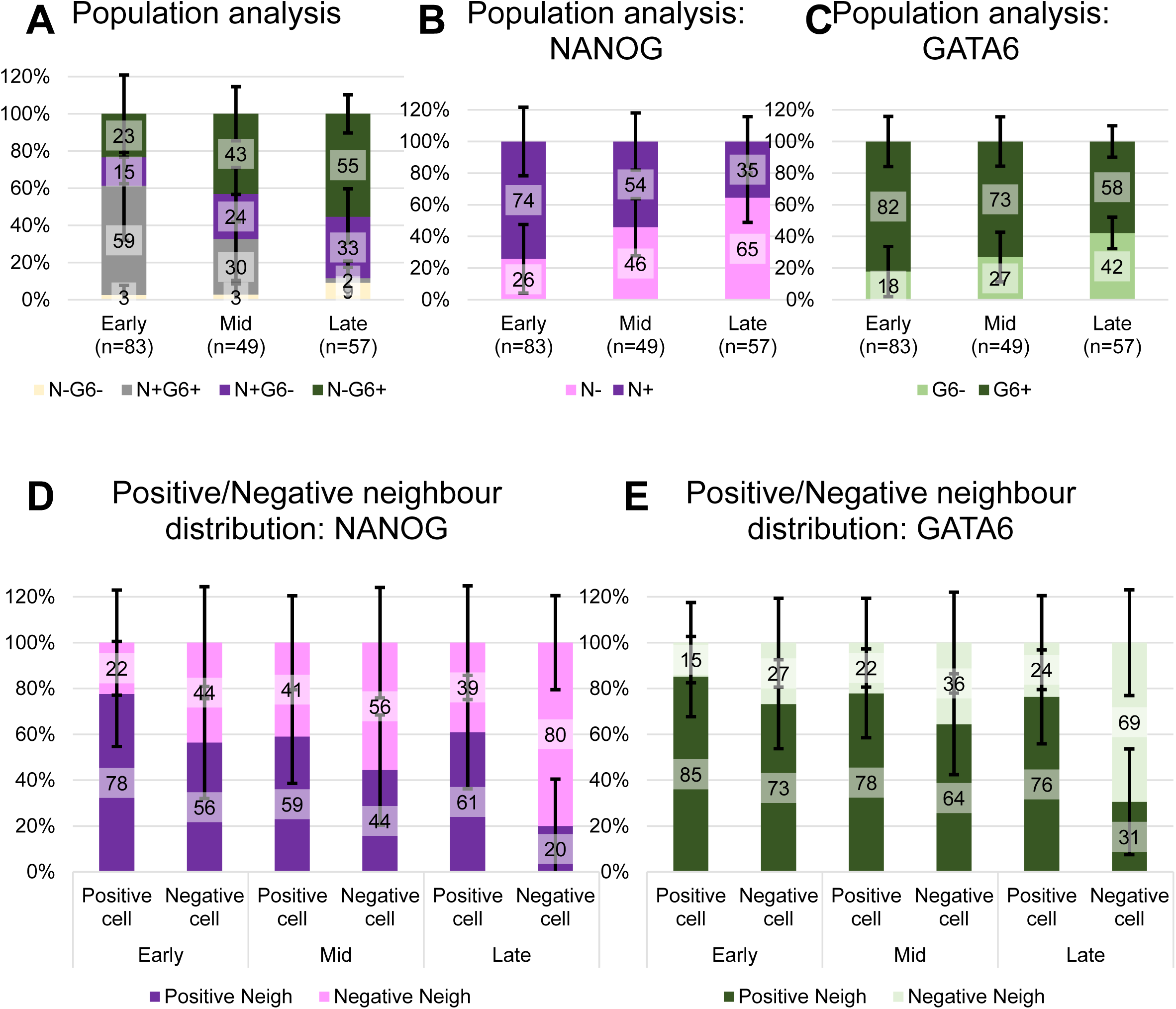
Average population and neighbourhood composition of all embryos in data set VI^20^. **(A)** Results of ICM population analysis indicating the average percentage of N-G6-, N+G6+, N+G6- and N-G6+ cells in early (n=83), mid (n=49) and late (n=57) blastocysts. Error bars show the standard deviation, here and in subsequent graphs. **(B-C)** Results of ICM population analysis indicating the average percentage of N+ or N-cells (B) and G6+ or G6-cells (C) in early, mid and late blastocysts. **(D-E)** Results of neighbour composition analyses for early, mid and late blastocysts indicating the average percentage of N+ or N-neighbours (D) and G6+ or G6-neighbours(E) of each ICM cell type.

**Supplemental Figure 7:**
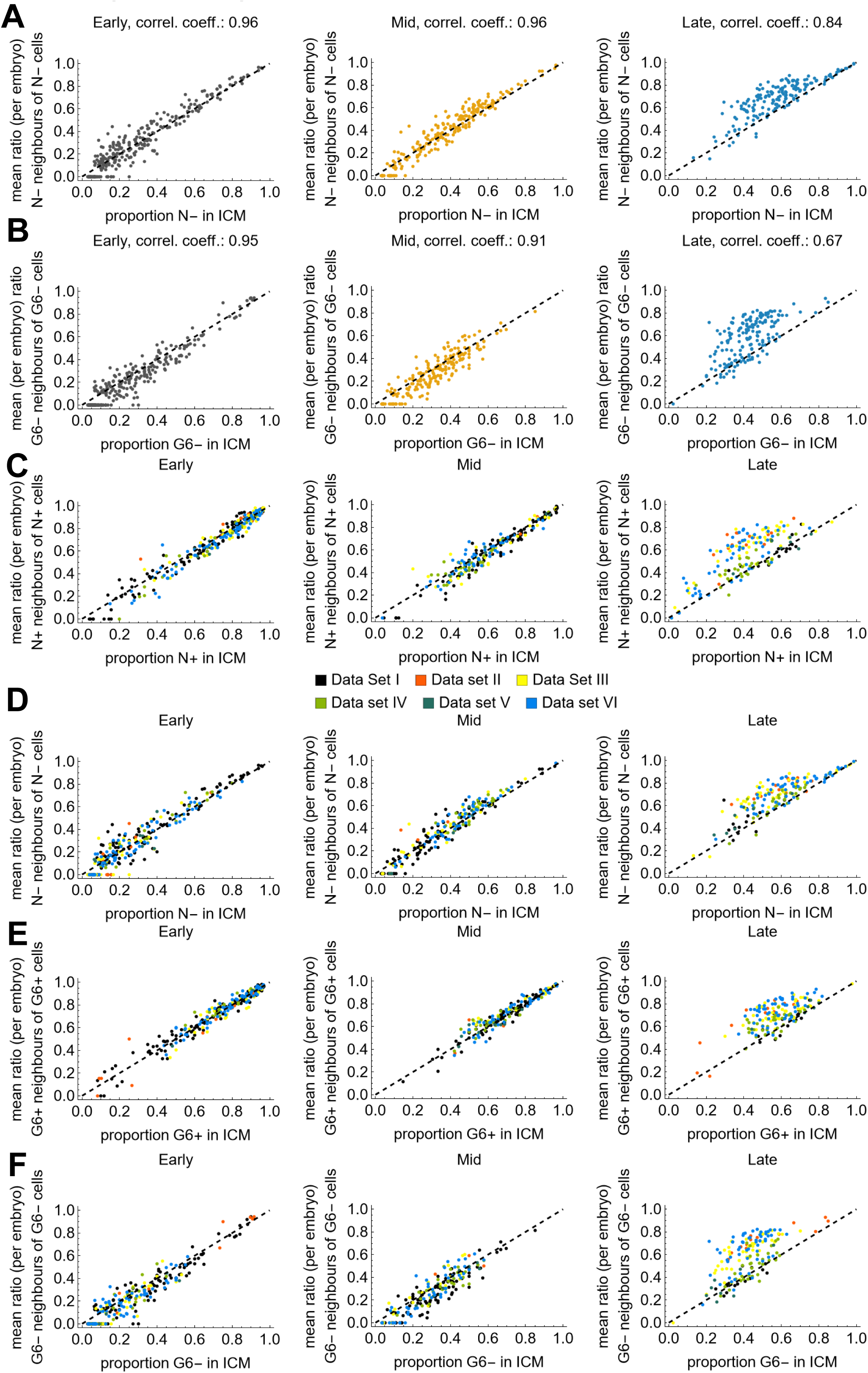
Single embryo neighbourhood quantification. **(A-B)** ICM composition scatter plot showing the mean ratio of N-neighbours of N-cells against the proportion of N- in each ICM (A) and the mean ratio of G6-neighbours of G6-cells against the proportion of G6-in each ICM against (B) in early (left), mid (middle) and late (right) blastocysts. Each dot represents one embryo. Pearson’s correlation coefficient is indicated above each plot. **(C-F)** ICM composition scatter plot showing the mean ratio of the indicated neighbour type of indicated cell types against the proportion of that cell type in each ICM in early (left), mid (middle) and late (right) blastocysts. The different colours indicate the individual data sets.

**Supplemental Figure 8:**
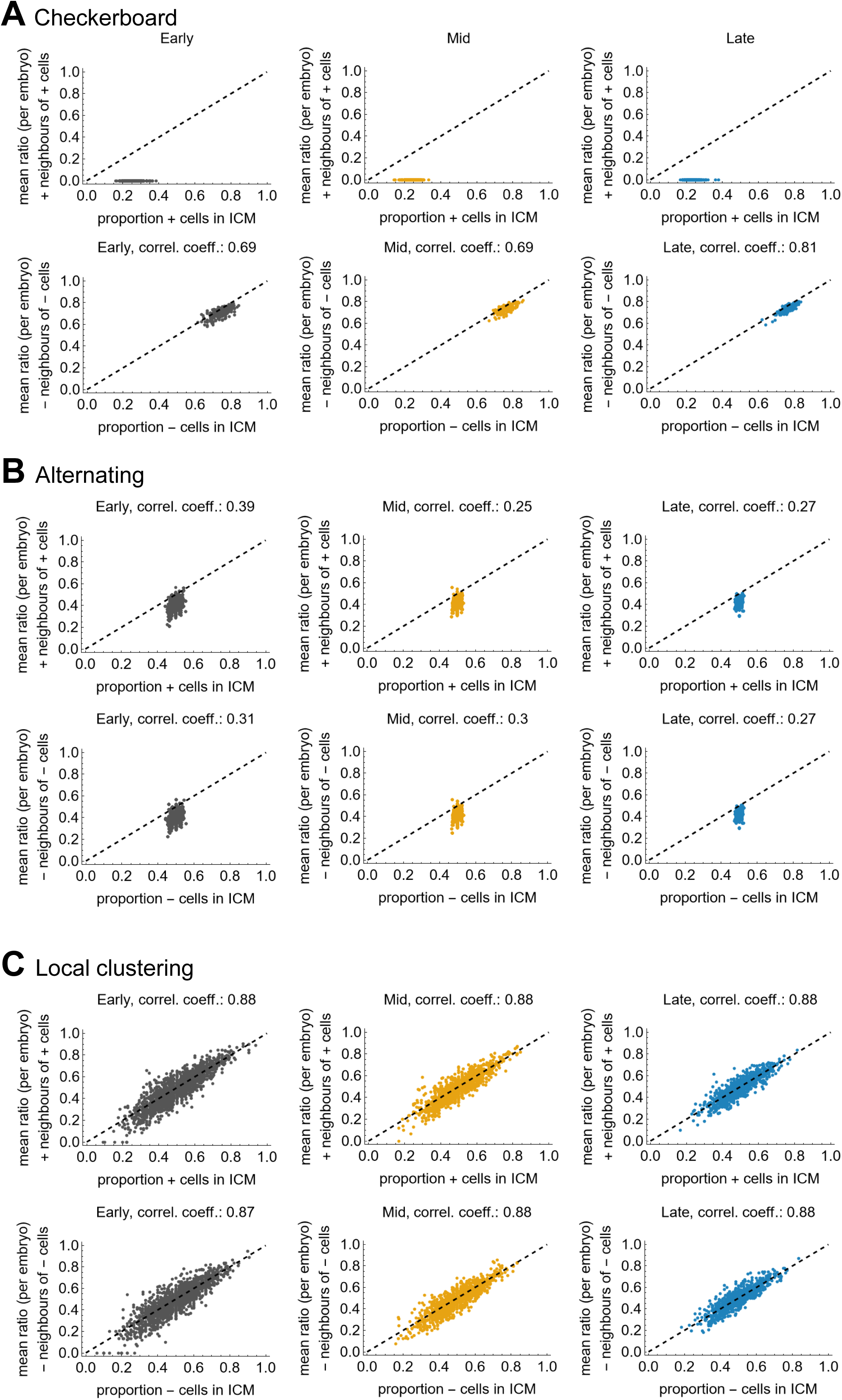
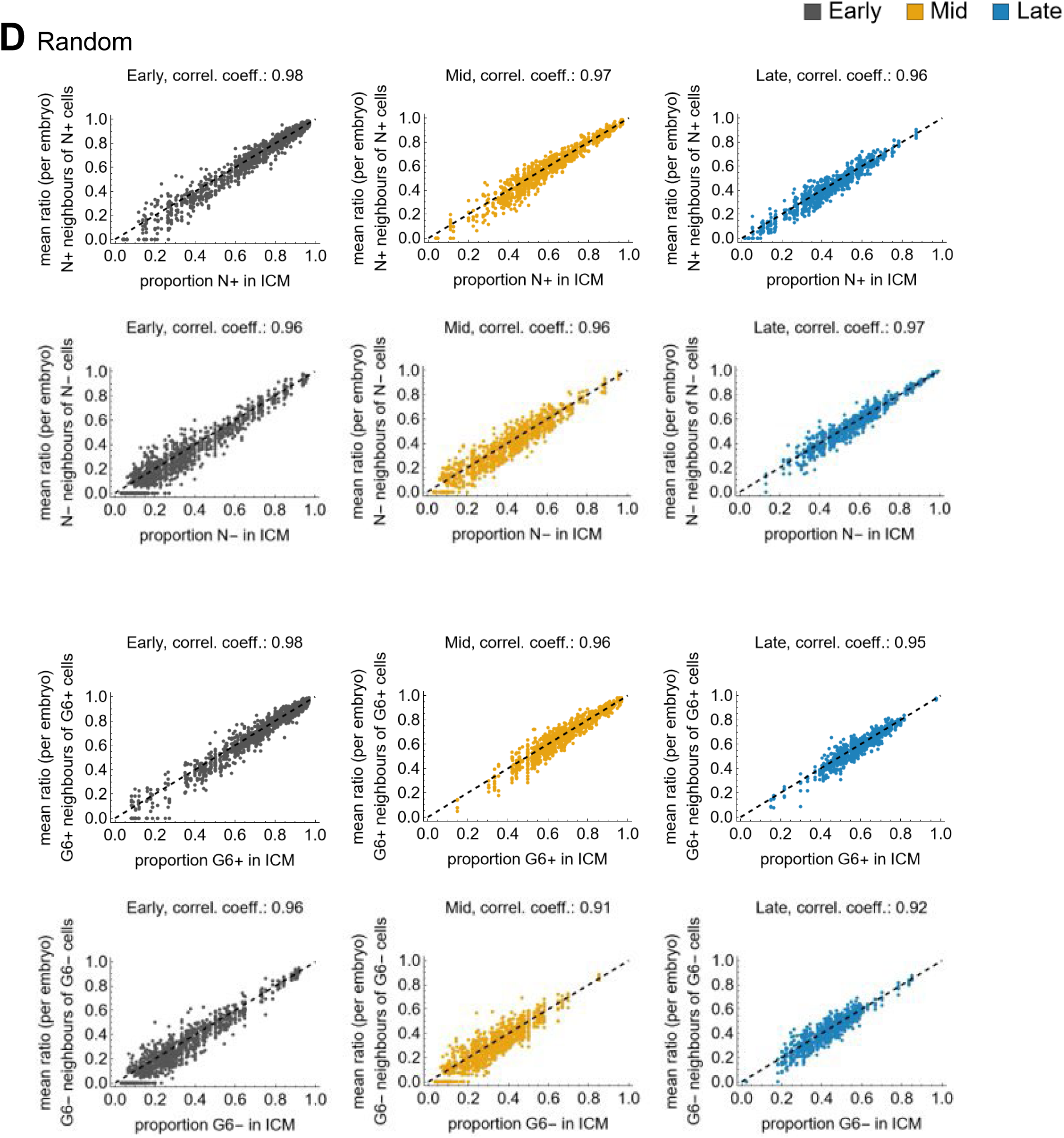
Single embryo neighbourhood for rule-based models. **(A-D)** ICM composition scatter plot for the simulation results. Scatter plots for the checkerboard (A), alternating (B), local clustering (C) and random (D) patterns showing the mean ratio of positive neighbours of positive cells against the proportion of positive cells in each modelled ICM in early (left), mid (middle) and late (left) modelled ICMs in each top row Each bottom row shows the analogous results for negative cells. Each dot represents one simulation for one ICM. Pearson’s correlation coefficient is indicated above each plot. Please note that for the neighbourhood composition of positive cells in the checkerboard pattern, Pearson’s correlation coefficient is not defined.

**Supplemental Figure 9:**
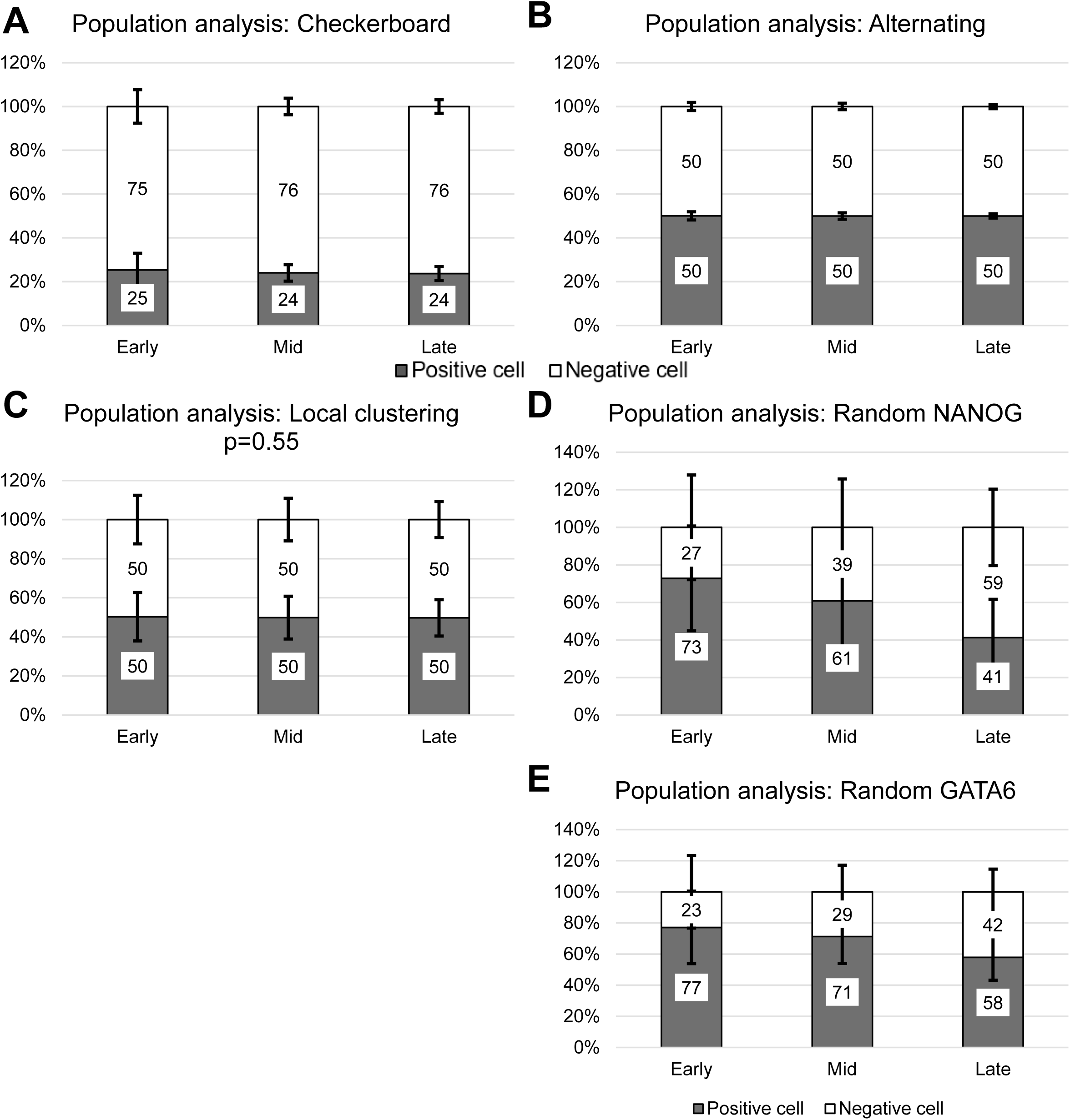
Average population and neighbourhood composition for rule-based models. **(A-E)** ICM population analysis of the checkerboard (A), alternating (B), local clustering with p=0.55 (C) and random patterns using NANOG data (D) or GATA6 data (E). Indicated is the average percentage of positive and negative cells in early, mid and late modelled ICMs.

**Supplemental Figure 10:**
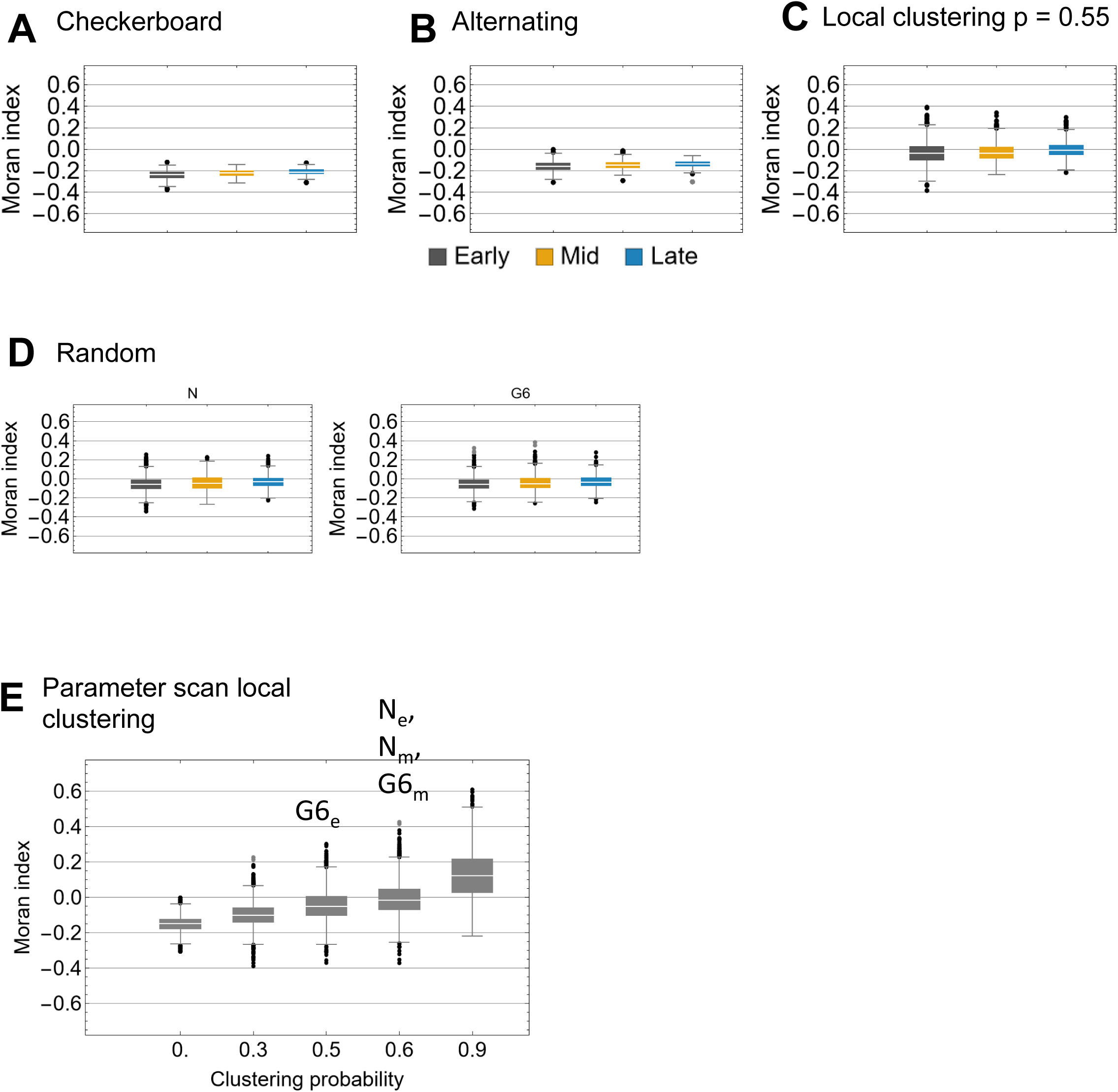
Morańs indices for rule-based models by stage. **(A-D)** Box-Whisker-Charts of Moran’s indices for the rule-based model patterns checkerboard (A), alternating (B), local clustering with p=0.55 (C) and random for NANOG data (D left) and GATA6 data (D right) for early (grey), mid (yellow) and late (blue) ICMs. **(E)** Box-Whisker-Charts of Moran’s indices of a parameter scan for the rule-based local clustering model patterns for the indicated clustering probabilities. To compare the modelling results with the experimental data, we performed a Welch’s t-test with Bonferroni correction with a significance level of 0.05. The letters above the boxes indicate for which experimental data, we did not detect a significant difference to the modelling results (G6_e_ and G6_m_ – GATA6 in early and mid blastocysts, respectively; N_e_ and N_m_ – NANOG in early and mid blastocysts, respectively)

**Supplemental Figure 11:**
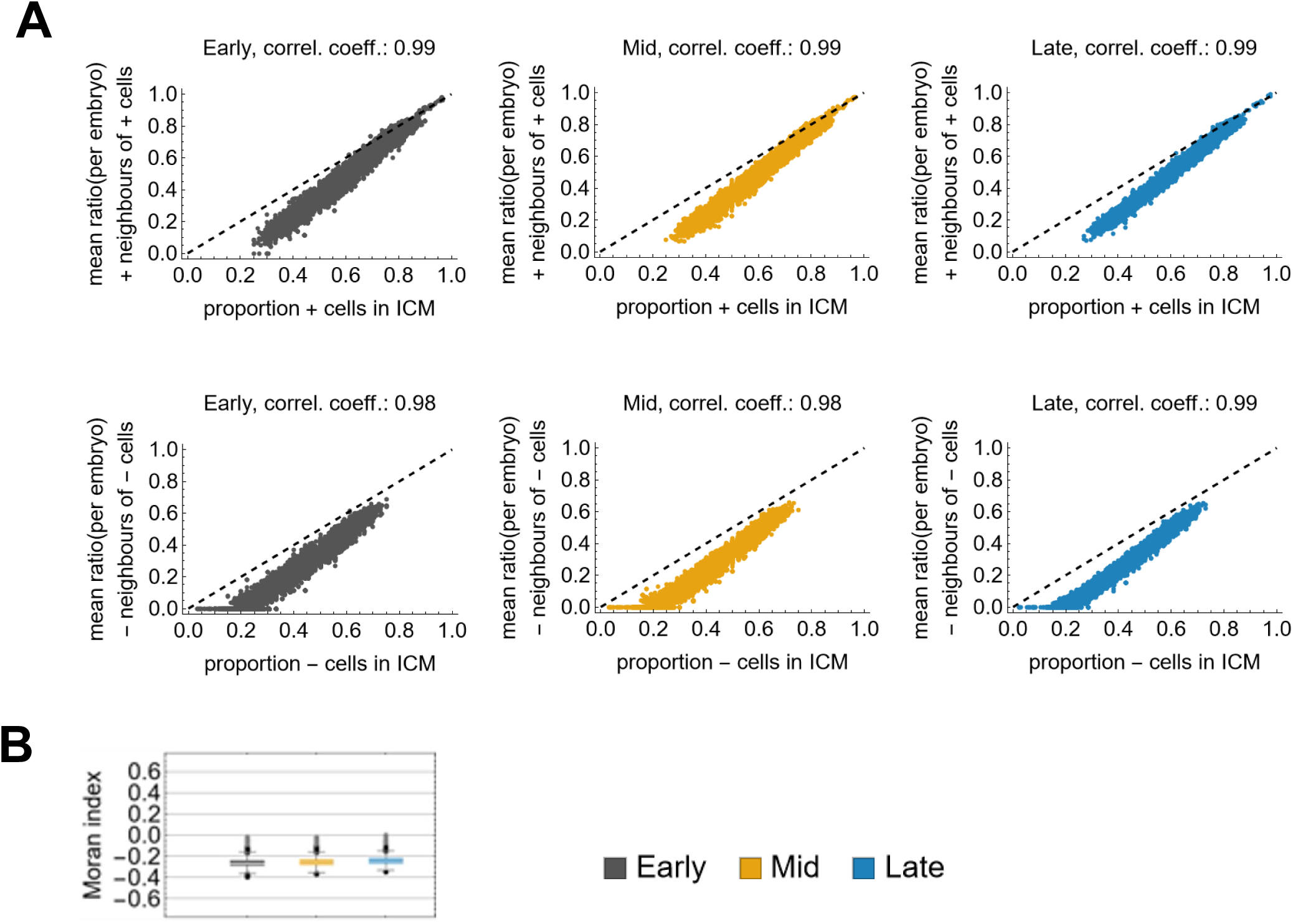
Nearest neighbour signalling: simulation results. **(A)** ICM composition scatter plot for the simulation results showing the mean ratio of positive neighbours of positive cells against the proportion of positive cells in each simulated ICM (top), and the mean ratio of negative neighbours of negative cells against the proportion of negative cells in each simulated ICM (bottom) in early (left), mid (middle) and late (left). Each dot represents one simulation for one ICM. Pearson’s correlation coefficient is indicated above each plot. **(B)** Box-Whisker-Charts of Moran’s indices for nearest neighbour simulations for early (grey), mid (yellow) and late (blue) ICMs.

**Supplemental Figure 12:**
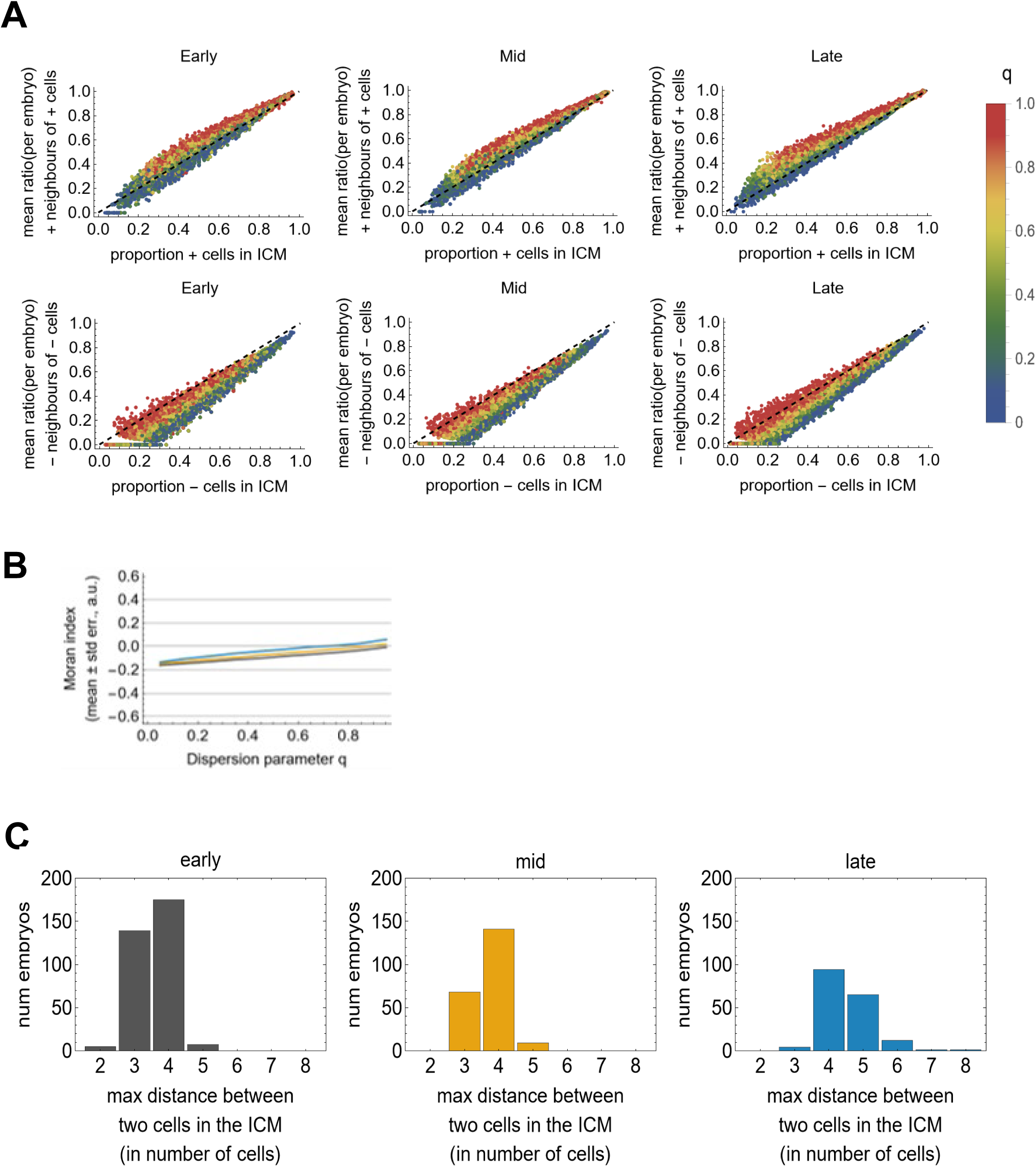
Distance-based neighbour signalling: simulation results. **(A)** ICM composition scatter plot for the simulation results showing the mean ratio of positive neighbours of positive cells against the proportion of positive cells for each simulation for an ICM (top), and the mean ratio of negative neighbours of negative cells against the proportion of negative cells for each simulation for an ICM (bottom) in early (left), mid (middle) and late (left). The colour code indicates the *q* values. Each dot represents one simulation for one ICM. **(B)** Moran’s indices against dispersion parameter q for simulations for early (grey), mid (yellow) and late (blue) ICMs. Each plot consists of a solid line for the average and a shaded area for the standard error of the mean (SEM). Please note that the SEM is very small and therefore hardly visible. **(C)** The distribution of maximum distances (in units of cells) between two cells in the ICM. The distances were calculated by the diameters of the cell graph representations of the ICMs.

**Supplemental Figure 13:**
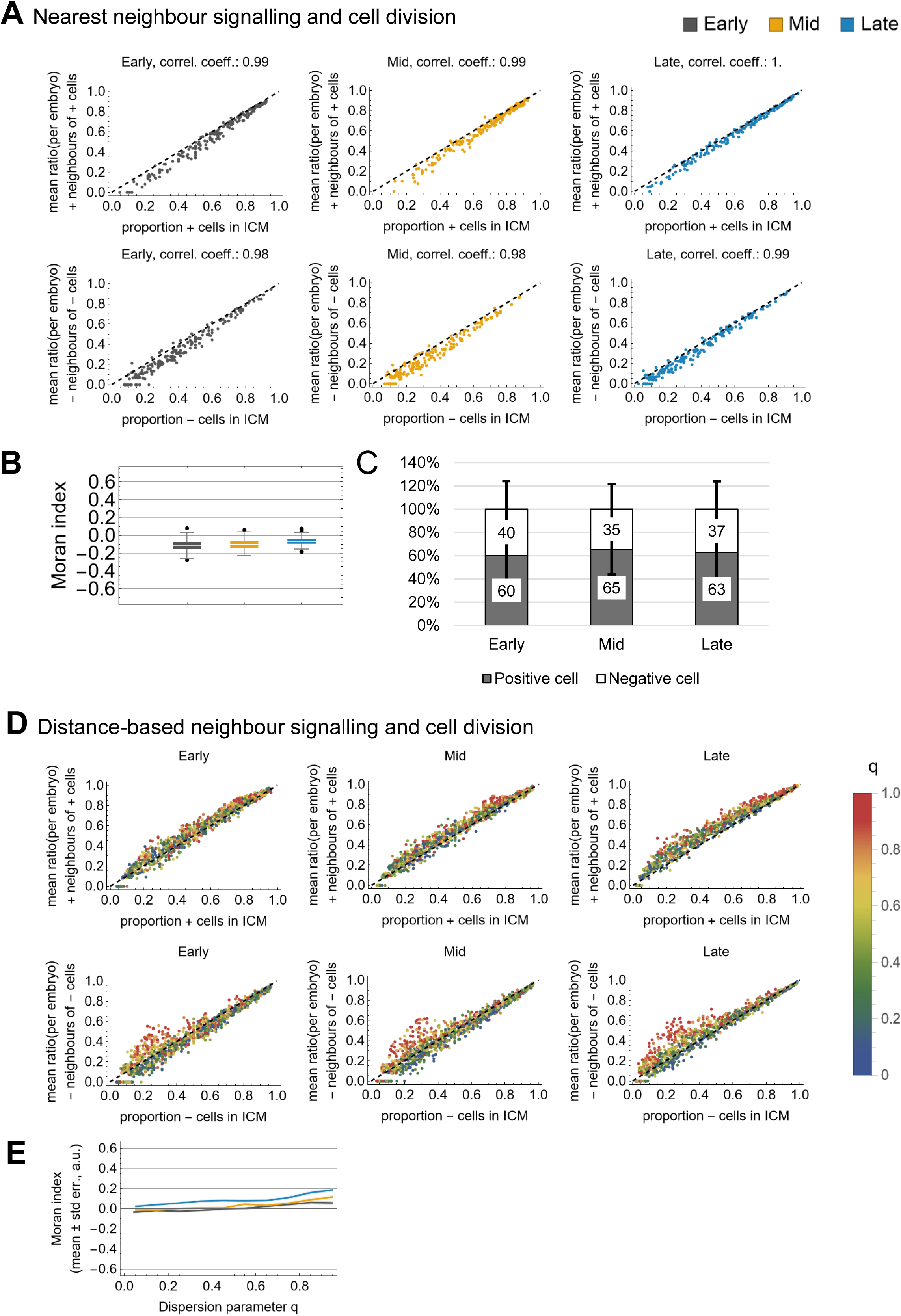
Nearest neighbour and distance-based neighbour signalling with cell division: Simulation results. **(A-B)** Nearest neighbour signalling with cell division: Simulation results (cont.). **(A)** ICM composition scatter plot for the simulation results showing the mean ratio of positive neighbours of positive cells against the proportion of positive cells (top) and negative neighbours of negative cells against the proportion of negative cells (bottom) in each simulated ICM at early (left), mid (mid), and late stage (right). **(B)** Box-Whisker-Charts of Moran’s indices for early (grey), mid (yellow) and late (blue) simulated ICMs. **(C)** ICM population analysis indicating the average percentage of positive and negative cells in the simulations for early, mid and late ICMs. **(D-E)** Distance-based neighbour signalling with cell division: Simulation results (cont.). **(D)** ICM composition scatter plot for the simulation results showing the mean ratio of positive neighbours of positive cells against the proportion of positive cells (top) and negative neighbours of negative cells against the proportion of negative cells (bottom) in each simulated ICM at early (left), mid (mid), and late stage (right). The colour coding indicates the *q* values. Each dot represents one simulation for one ICM. **(E)** Moran index against dispersion parameter *q* for simulations for early (grey), mid (yellow) and late (blue) ICMs. Each plot consists of a solid line for the average and a shaded area for the standard error of the mean (SEM). Please note that the SEM is very small and therefore hardly visible.

**Supplemental Figure 14:**
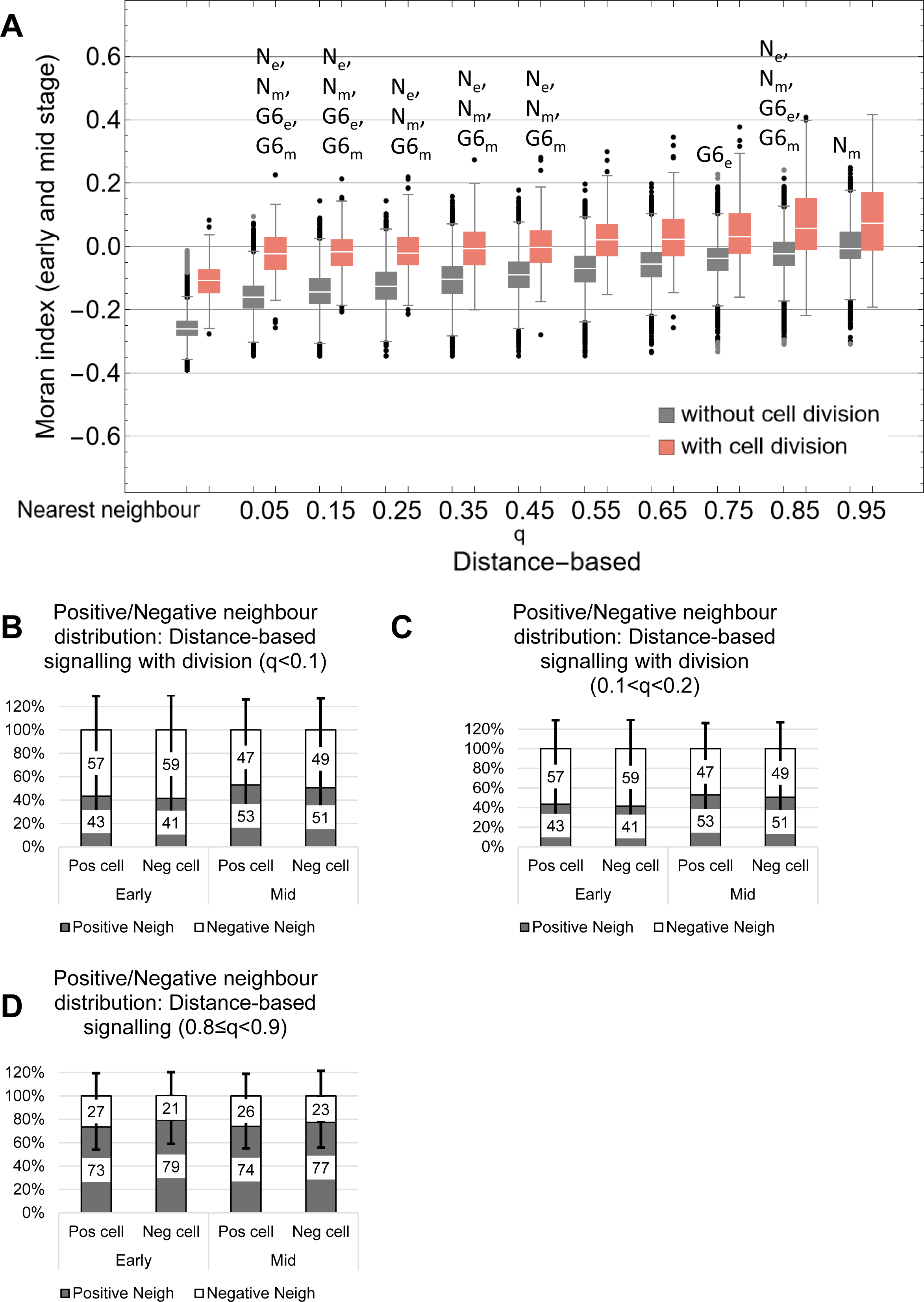
Signalling models with experimental data comparisons. **(A)** Box-Whisker-Charts of Moran’s indices for nearest neighbour and distance-based neighbour signalling without (grey) and with (red) cell division. The distance-based neighbour signalling models were collected in bins of width 0.1 based on the respective q value. The letters Ne, Nm, G6e and G6m refer to the experimental data for NANOG in early, NANOG in mid, GATA6 in early and GATA6 in mid stage blastocysts. We performed a Welch’s t test with Bonferroni correction and a significance level of 0.05 to compare the different model results to the experimental data. The letters indicate those comparisons for which we could not detect a significant difference. **(B)** Neighbour composition analyses for the simulation results for distance-based signalling model with *q*<0.1 and cell division in early, mid and late blastocysts indicating the average percentage of positive or negative neighbours of each ICM cell type. **(C)** Neighbour composition analyses for the simulation results for distance-based signalling model with 0.1<*q*<0.2 and cell division in early, mid and late blastocysts indicating the average percentage of positive or negative neighbours of each ICM cell type. **(D)** Neighbour composition analyses for the simulation results for distance-based signalling model with 0.8≤*q*<0.9 and no cell division in early, mid and late blastocysts indicating the average percentage of positive or negative neighbours of each ICM cell type.

## STAR Methods

### RESOURCE AVAILABILITY

#### Lead contact

Further information and requests for original images and data should be directed to and will be fulfilled by the lead contact, Silvia Muñoz-Descalzo. Enquiries about models and code should be directed to Sabine C. Fischer

#### Materials availability

This study did not generate new unique reagents.

#### Data and code availability

All data analysis was done in Mathematica 13.0. The model simulations were done in Python 3.7 using Jupyter notebooks as an interactive development environment. All processed data as well as the code used to transform and analyse data is available at https://github.com/scfischer/fischer-et-al-2023 archived at Zenodo https://doi.org/10.5281/zenodo.7867284.

The following dataset was generated:

| Authors | Publication year | Data set title | Data set URL |
| --- | --- | --- | --- |
| Fischer SC, Schardt S, Lilao-Garzon J, Muñoz-Descalzo S | 2023 | The salt-and-pepper pattern in mouse blastocysts is compatible with signalling beyond the nearest neighbours | https://doi.org/10.5281/zenodo.7867284 |

The following previously published datasets were used:

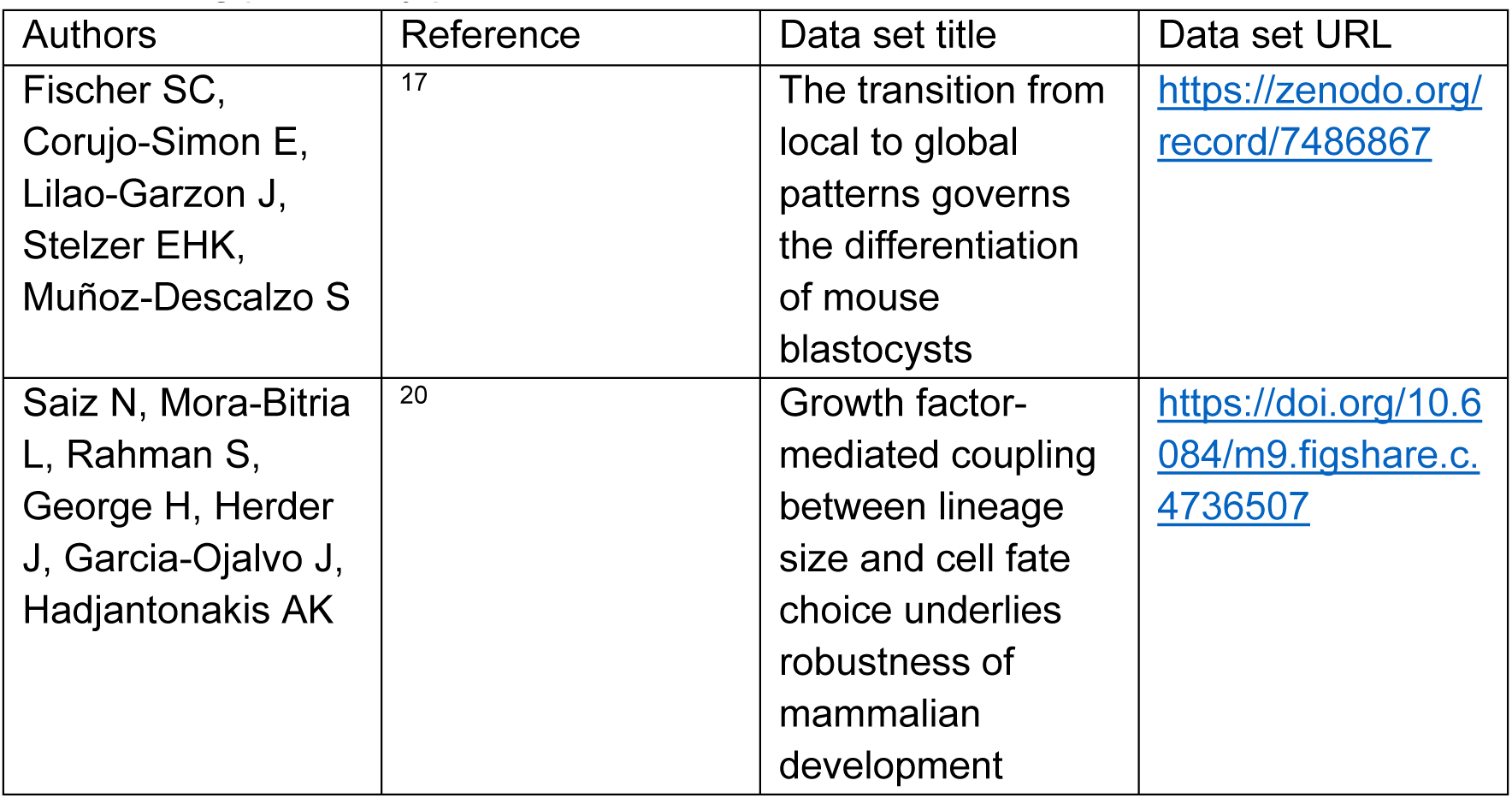
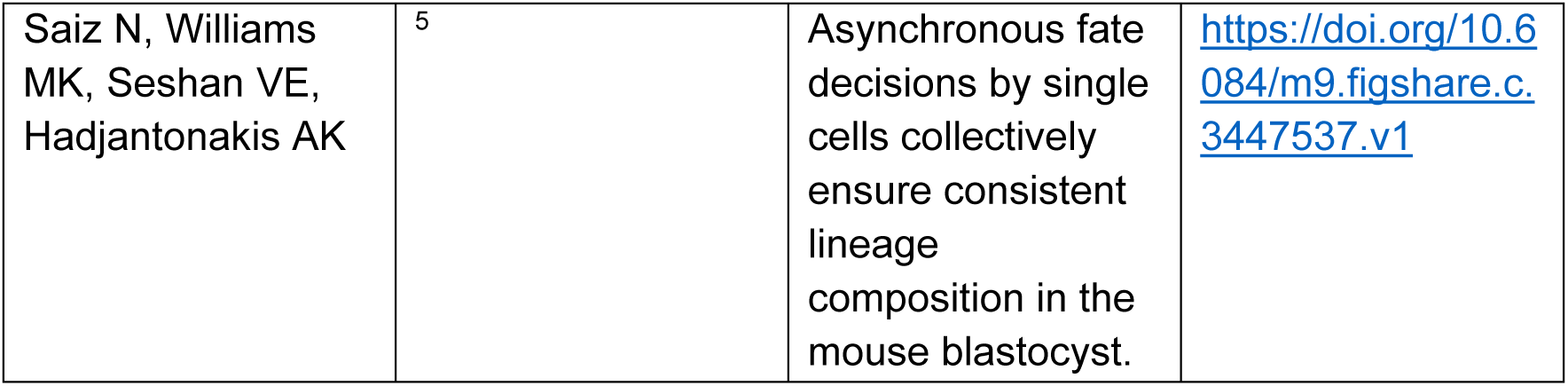

### EXPERIMENTAL MODEL AND SUBJECT DETAILS

SWISS mice were bred in house at the Universidad de Las Palmas de Gran Canaria (ULPGC) at the Institute for Biomedical and Healthcare Research (IUIBS) animal facility. All mice were housed with controlled room temperature (20-24°C) and relative humidity (55-72%), and a 12-h light-darkness cycle. All animal studies were conducted following National and European regulations (RD 1201/2005, Law 32/2007, EU Directive 2010/63/EU). The sex of embryos was not determined for the analyses conducted in this study. Embryos were classified, according to their total cell number, into early (32-64 cells), mid (65-90 cells) and late stage (more than 90 cells).

## METHOD DETAILS

### Experimental Data

#### Data sets

In this work, we generated one new data set and reanalysed three existing ones. Two of the data sets were imaged in our lab and the other two were obtained from publications from the Hadjantonakis’ lab (Sloan Kettering Institute). For our analysis, we require for each cell the information whether it is a TE or an ICM cell, its centroid position and its expression levels of NANOG and GATA6. The cells need to be classified as NANOG+ or NANOG-as well as GATA6+ or GATA6-. Furthermore, we need a list of cells that are direct neighbours to the given cell. This happened in four steps: Imaging, Image analysis, population assignment and neighbourhood extraction. For most data sets, the first three steps were performed and described in previous publications (Table 1). The neighbourhood extraction for all data sets was done in this study.

**Table 1:**
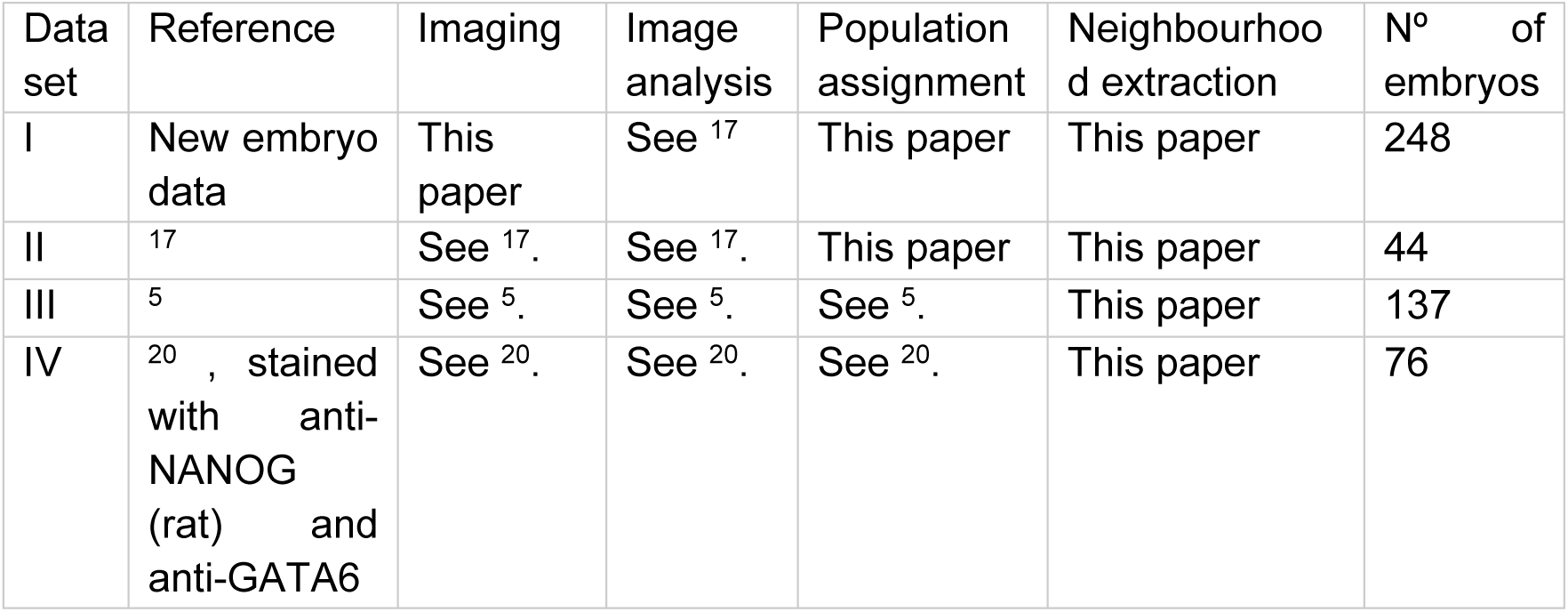
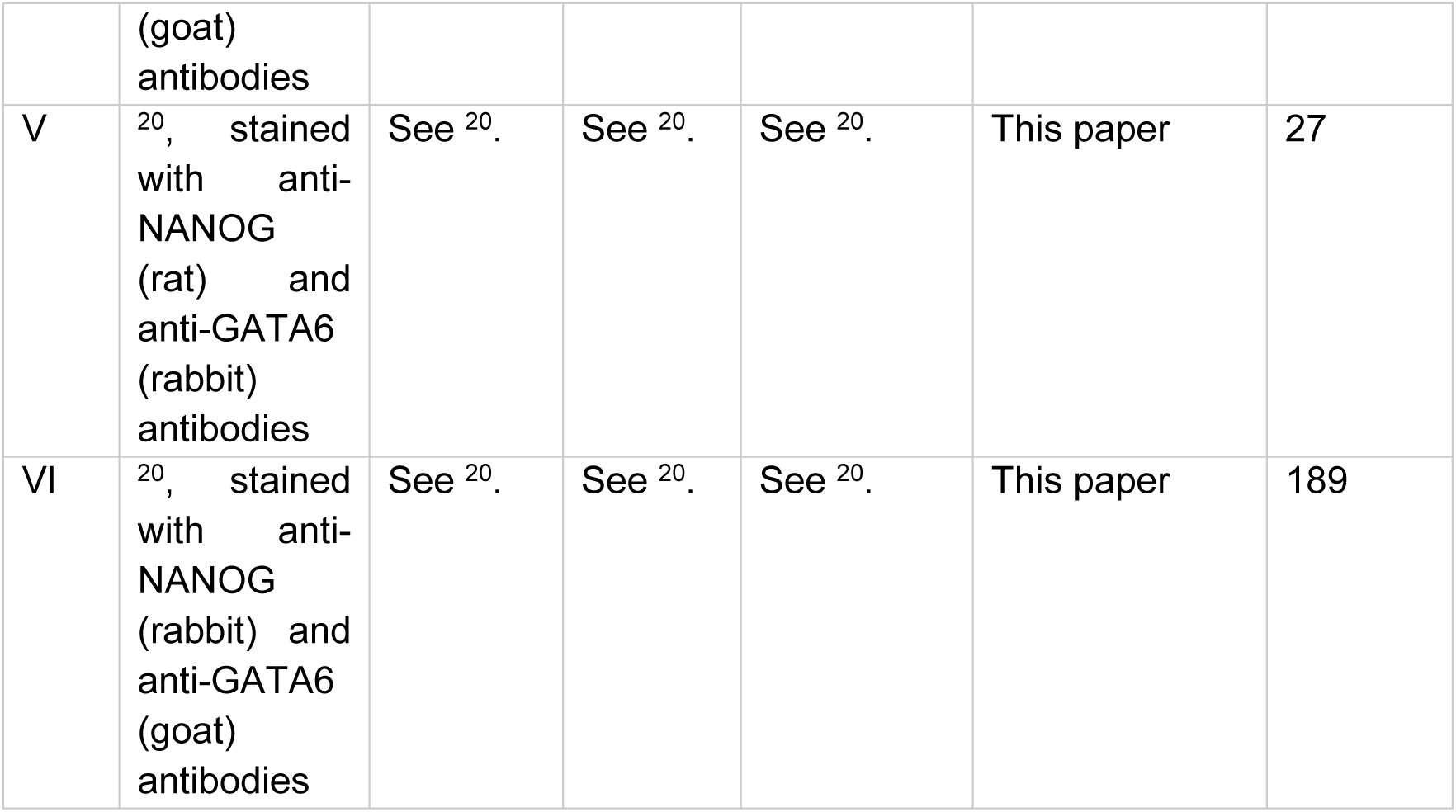
Overview of data sets and processing steps

#### Embryo Collection

Wild-type Swiss embryos were generated by in-house breeding and natural mating. Detection of copulation plug confirmed successful mating; the resulting embryos were then considered embryonic day (E) 0.5. Embryos were isolated in M2 medium (Embryomax®).

#### Immunofluorescence

Embryos were prepared for immunofluorescence as previously described ^50^. *Zona pellucida* was removed using acid Tyrode’s. Embryos were fixed in 4% paraformaldehyde in PBS for 15 minutes, then rinsed in PBS containing 3 mg/ml polyvinylpyrolidone (PBS/PVP), permeabilised in PBS/PVP containing 0.25% Triton X-100 for 30 minutes and blocked in blocking buffer, which comprised PBS containing 0.1% BSA, 0.01% Tween 20 and 2% donkey serum. Antibodies were diluted in blocking buffer and embryos incubated at 4°C overnight. Primary antibodies were anti-NANOG (eBioscience, 1:200), anti-GATA6 (R&D Systems, 1:200) They were rinsed three time in blocking buffer for 15 minutes each and incubated in secondary antibody solution for 1 hour in the dark. Secondary antibodies labelled with Alexa fluorophores (Invitrogen) were diluted 1:1000 in blocking buffer. Embryos were then rinsed three times in blocking buffer, incubated briefly in increasing concentrations of Vectashield before mounting on glass slides with Vaseline bridges to prevent their crushing in small drops of concentrated Vectashield, and subsequently sealed with nail varnish. DAPI (1:1000) was used to stain the nuclei.

#### Imaging and automated image analysis for data set I

Embryos were imaged using a Zeiss LSM Zeiss LSM700 and a Plan-Apochromat 40x/1.3 Oil DIC (UV) VIS-IR M27 objective, with optical section thickness of 1 μm. Embryo images were processed and analysed as previously described ^17^. All images in each imaging session were obtained using the sequential scanning mode, with the same conditions of laser intensity, gain, and pinhole, and were processed in exactly the same way. The range indicator palette option (Zeiss AIM/ZEN software) was used to ensure that no oversaturated images were taken. The three-dimensional image stacks were segmented using MINS ^51^. Cells were automatically assigned to ICM or TE. The features of the cell nuclei were extracted including the nuclear centroid and volume, together with the mean intensity of NANOG and GATA6 for each nucleus. The automatically assigned TE or ICM fate was manually checked using acquired images and ImageJ. Extreme errors (over-segmentation and pyknotic nuclei) in the segmentation were removed manually when correcting the classification of TE versus ICM.

#### Population assignment for data sets I and II

Depending on the thickness of a sample, confocal imaging results in fluorescence intensity decay along the z-axis. We quantified this decay for each embryo by fitting a linear regression model to the expression values of the trophectoderm cells. The slope of the regression curve then provided the factor to correct for the decay in the ICM cells.

In our setup, we mount the embryos, which results in a slight squeezing along the z-axis of the image. The correction for squeezing was performed as previously described ^17^.

To determine the thresholds between high and low expression levels of NANOG and GATA6, we performed a mixture analysis with PAST ^52^ for each transcription factor and imaging session independently.

#### Neighbourhood extraction for all data sets I-VI

For the neighbourhood analyses we needed to determine the direct neighbours of each cell based on the centroid positions. To this end, we derived a cell graph representation for each embryo in all data sets as described previously ^17, 53^ and extracted a list of direct neighbours for each cell.

### Modelling and simulations

To characterise the salt-and-pepper pattern, we generated artificial patterns and compared their properties to those of the experimental data. Our models range from simple rule-based models to more sophisticated approaches that include nearest neighbour or distance-based neighbour signalling.

#### Rule-based models

We used the nuclei centroids and cell graphs from the experimental data of all data sets and all stages and assigned positive or negative fate to each cell. For each ICM, we generated patterns based on the following rules:

- *Checkerboard:* One cell is assigned a positive fate. All direct neighbours of that cell are assigned a negative fate. The next cell in the list of centroids that has not been assigned a fate yet obtains a positive fate. This procedure is repeated until all cells have a positive or negative fate. In the resulting pattern, positive cells have only negative neighbours.
- *Alternating:* Starting with one centroid, a list of centroids is generated such that the next one is the closest centroid in the ICM to the previous one with respect to Euclidean distance. The cells in this list are assigned alternating positive and negative fate. This results in a pattern in which cell fate is alternating between nearest neighbour cells, i.e. if a cell is positive the cell that is closest with respect to Euclidean distance is negative and so on.
- *Local cluster with probability p*: As for the alternating pattern, we start with one centroid and generate a list of centroids in which the next one is the closest centroid in the ICM to the previous one with respect to Euclidean distance. The first cell in the list is randomly assigned a positive or negative state. The next entry in the list, i.e. the closest cell based on Euclidean distance, is assigned the same state as the previous cell with a given probability p. We analysed modelling patterns for p=0, 0.1, 0.2,…, 0.9 and compared their Moran’s indices to those of the experimental data (Sup Fig. 10). We found the patterns for p = 0.5 and p = 0.6 to both fit some of the experimental conditions. Therefore, we further investigated the pattern for p = 0.55 and found it to fit the experimental data for early and mid embryos sufficiently well based on hypothesis testing (see below)
- *Random:* We extract the number of positive and negative cells from the experimental data for the given ICM. Then we randomly assign that number of positive and negative fate to the cells. This procedure is conducted both for NANOG and for GATA6, resulting in two types of patterns.

For the checkerboard rule, we did not expect the choice of the first cell to create differences in the neighbourhood properties of the pattern. Therefore, our implementation was deterministic, and we generated only one replicate per embryo. For the alternating, the local cluster and the random rule, we generated five replicates for each ICM.

#### Signalling models

First, we implemented a model for neighbour signalling on a fixed tissue. We used the embryo geometries including cell number and cell centroids from the experimental data of all data sets I-VI as the underlying tissue geometries and employed our model for transcriptional regulation based on intra- and intercellular signalling of two transcription factors u and v ^27^. This resulted in three-dimensional cell fate patterns of u-v+ and u+v-, i.e. positive and negative cells. We performed 100 replicates per embryo geometry for each signalling model: nearest neighbour and distance-based. For the distance-based neighbour signalling, we further varied the dispersion parameter *q*. We chose a different value for each simulation randomly from a uniform distribution on (0,1).

As a second approach, we conducted simulations on a growing tissue. We started with a single cell of radius 0.75 and let the tissue evolve while the interactions for the gene regulatory network took place. Hence, each simulation time step comprised four distinct processes:

- Cell growth
- Cell division
- Adhesion and repulsion
- Transcriptional regulation

In this case, we could not use the embryo geometries from the experimental data. Instead, we determined the average ICM cell counts of early, mid and late blastocysts as 22, 25 and 42 cells, and chose the final times for our simulations accordingly.

For the nearest neighbour signalling, we performed 200 simulations for each stage. For the distance-based neighbour signalling, we performed 1200 simulations for each stage and chose for each simulation a different value for the dispersion parameter *q* randomly from a uniform distribution on (0,1).

A detailed description of the modelling approaches can be found in the supplementary text.

### QUANTIFICATION AND STATISTICAL ANALYSES

**Moran’s index** is a measure to quantify spatial autocorrelations. Its original form was first introduced in ^23^ to analyse the distribution of soil fertility over a field. For details, please refer to the supplementary text.

In the **Box-Whisker-Chart**, the boxes denote the range from 25% to 75% quantile. The white line indicates the median. Values that are larger than the 75% quantile plus 1.5 times the interquartile range are near outliers, marked in black. Values that are larger than the 75% quantile plus 3 times the interquartile range are far outliers, marked in grey.

ICMs that contain only one cell type were excluded from the analysis. We excluded 51 early, 2 mid and 2 late embryos for NANOG, and 64 early, 19 mid and 6 late embryos for GATA6.

For the simulation results of the distance-based neighbour signalling model, we displayed Moran’s index as a function of q. To this end, results were binned according to their q values in bins of size 0.1.

To determine the best fitting model, we compared the model and simulations results with the experimental results both visually and by employing **hypothesis testing**. Unless otherwise stated, we used Welch’s t-test with Bonferroni correction and a significance level of 0.05.

## ADDITIONAL RESOURCES

### KEY RESOURCES TABLE

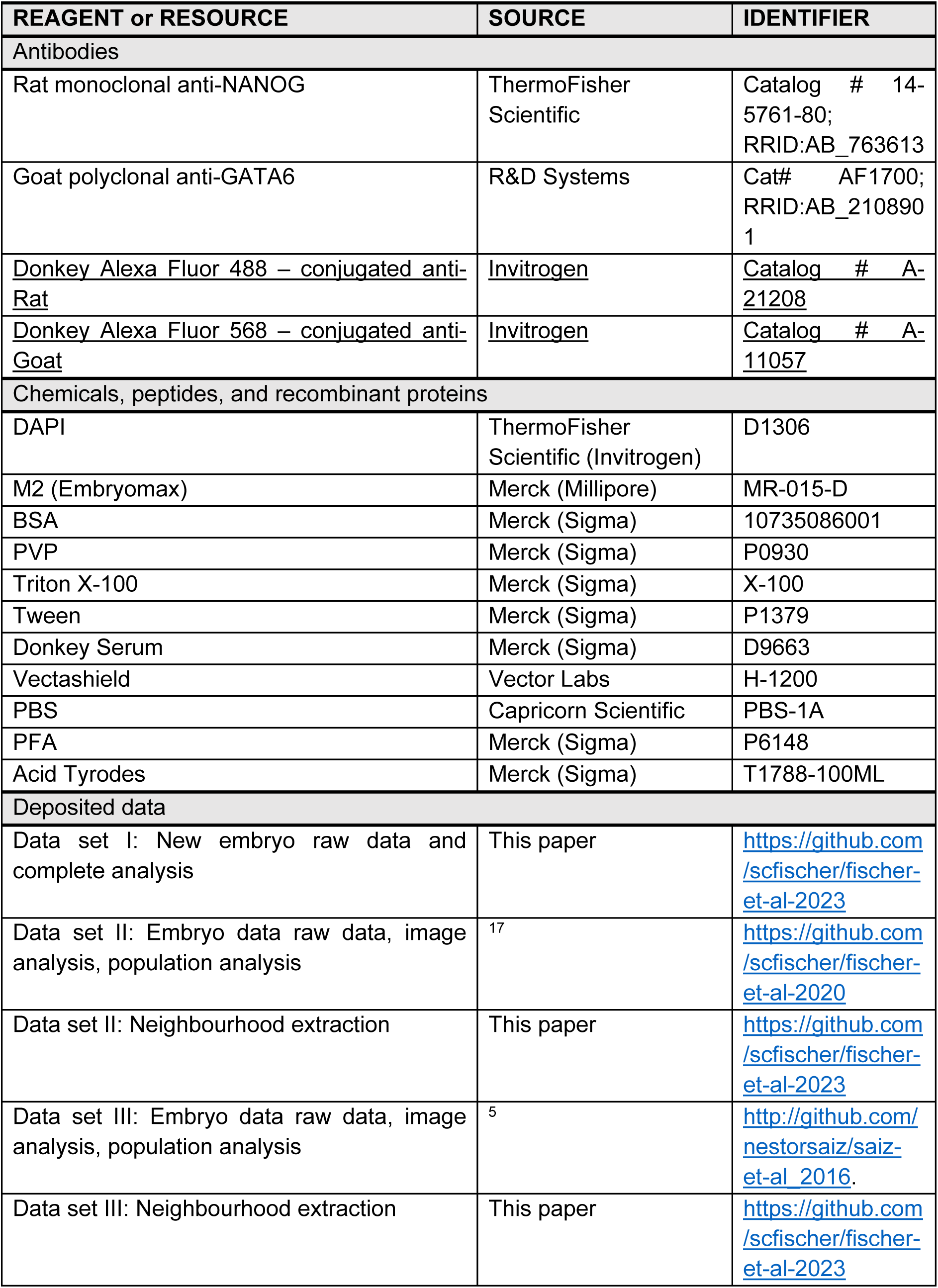
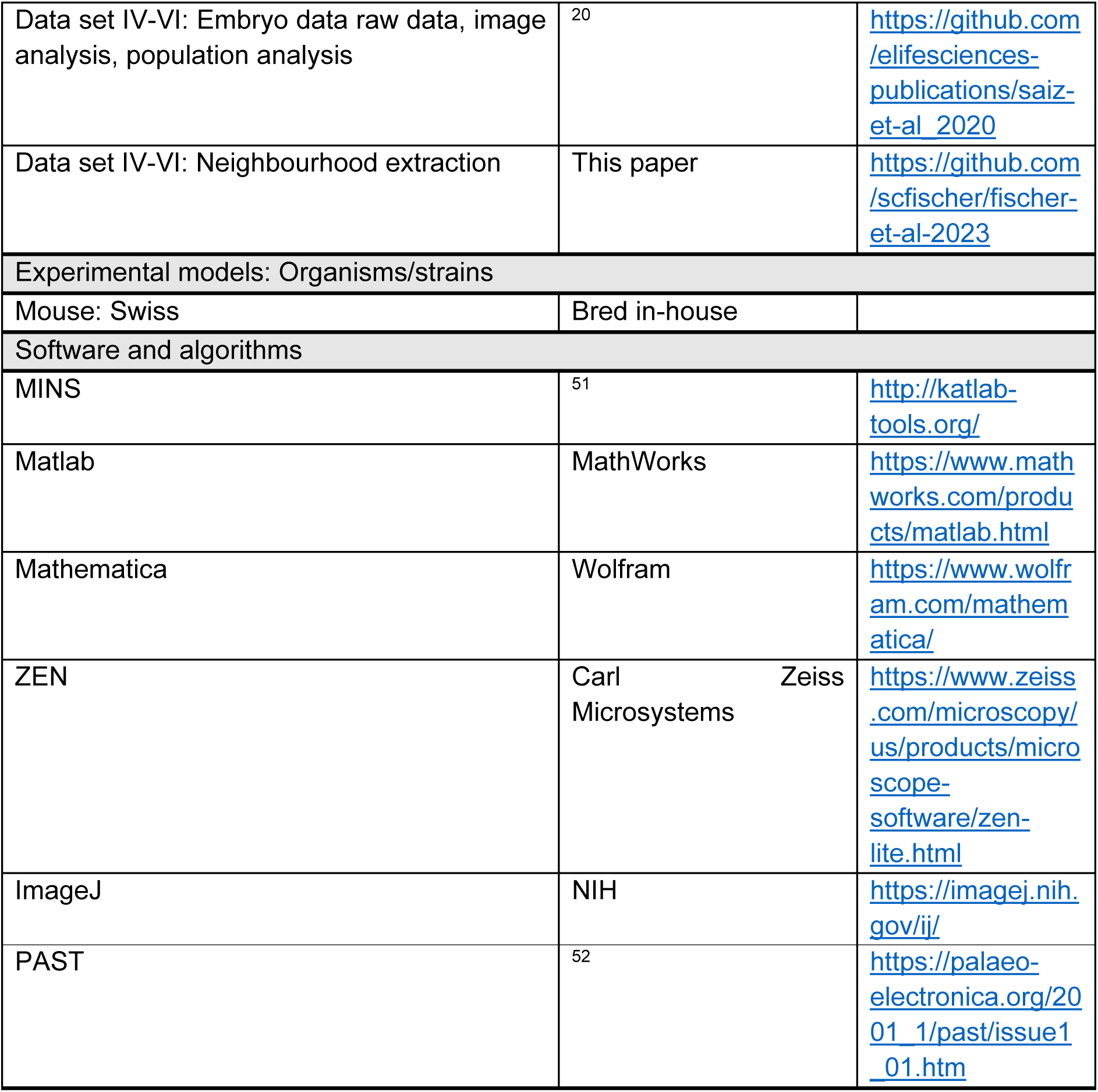

