## Supplemental details for "The salt-and-pepper pattern in mouse blastocysts is compatible with signalling beyond the nearest neighbours"

### 1 Measuring spatial autocorrelation

To quantify spatial autocorrelation in the cell fate patterns, we used Moran's index. For a variable of interest $x \in \mathbb{R}^n$, it is defines as

$$I := \frac{n}{\sum_{i,j} a_{i,j}} \frac{(x-\bar{x})^T A(x-\bar{x})}{(x-\bar{x})^T (x-\bar{x})},$$

where $\bar{x}$ denotes the mean of $x$ and $A$ is a matrix representation of the connectivity of the $n$ cells in a tissue, here the ICM. We chose $A$ as the adjacency matrix of our Delaunay cell graph. The fates of the cells are represented by $x$. Considering that we only incorporate two cell fates in our study, namely positive and negative, $x$ is given by a vector with entries $x_i \in 0, 1$ for $i = 1, \ldots, n$. The value of $I$ is independent on which value is assigned to which fate.

To get an intuition of Moran's index as a measure of spatial patterns, we consider two extreme cases in a synthetic two-dimensional example on an 8 x 8 grid (Fig I). In a checkerboard pattern, equal cells are never adjacent to each other as long as the number of neighbors does not exceed four. Thus, there is no auto-correlation between the cells leading to the minimal value $I = -1$. In the case of spatial separation, cells are mostly neighbours to equal cell types resulting in values of $I$ close to 1.

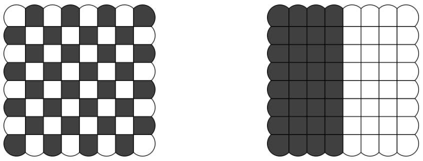

(a) $I = -1$ (b) $I = 0.857$

Figure I: Illustration of Moran's index on a two-dimensional 8 x 8 grid.

### 2 Agent-based modelling of cell differentiation in mouse ICM

The full computational model underlying the results in Fig 8 in the main text follows a time resolved process comprised of four distinct processes in every time step. These are:

- Cell growth
- Cell division
- Adhesion and repulsion
- Compute cell graph
- Transcriptional regulation

In Figs 6 and 7 in the main text, we considered only transcriptional regulation on a fixed tissue geometry. In the following sections, we provide the details for the individual steps.

### 2.1 Cell growth

The best fitting mathematical description for cell growth is still under debate [1]. Potential mechanisms are linear [2] as well as exponential [3] cell growth. In addition, cell size control, limiting the maximum cell size [4], might be involved. More recently, it has been shown that the growth rates of cells vary between different stages of the cell cycle [5]. Due to the large number of unknowns and since the main focus of our model is not on the details of cell growth, we used a simplified approach to model the growth of individual cells. We considered that cells grow by absorbing nutrients in the surrounding medium. The larger a cell becomes, the more nutrients it will be able to absorb through its surfaces, which suggests an exponential growth. However, nutrients are used to sustain the cell's own metabolism which creates a limitation to cell size. A prominent model that captures such a behavior is the logistic growth model. We applied it to the cell radius $r$, i.e.

$$\frac{dr}{dt} = \lambda r(r^* - r), \tag{1}$$

where $\lambda$ denotes a constant growth rate. The solution to this equation for initial condition $r(t_0) = r_0$ is given by

$$r(t) = \frac{r^*}{1 + \left(\frac{r^* - r_0}{r_0}\right) e^{-\lambda r^*(t-t_0)}}, \tag{2}$$

### 2.2 Cell division

Cell division is usually described as a function of either the elapsed time [6] or the size of a cell [7]. In the latter case, cell division has been successfully implemented for in silico generation of mouse ICM organoids. The division itself has been described as a stochastic process of a cell's radius. Likewise, we used a stochastic approach towards cell division based on the cell's radius. We used the cumulative distribution function (CDF) $F(r)$ of a truncated normal distribution to define the probability that a cell will have divided up to some radius $r$. The CDF is defined via

$$\begin{aligned} F(r) &= \frac{\mathrm{erf}(\rho) - \mathrm{erf}(\rho_{\min})}{\mathrm{erf}(\rho_{\max}) - \mathrm{erf}(\rho_{\min})}, \qquad r \in [r_{\min}, r_{\max}], \\ \rho &= \frac{r - \mu_{\mathrm{div}}}{\sqrt{2}\sigma_{\mathrm{div}}}, \qquad \rho_{\min} = \frac{r_{\min} - \mu_{\mathrm{div}}}{\sqrt{2}\sigma_{\mathrm{div}}}, \qquad \rho_{\max} = \frac{r_{\max} - \mu_{\mathrm{div}}}{\sqrt{2}\sigma_{\mathrm{div}}}, \end{aligned} \tag{3}$$

where $\mu_{\mathrm{div}}$ and $\sigma_{\mathrm{div}}$ are the mean and standard deviation of this distribution. We chose $\mu_{\mathrm{div}} = \frac{1}{2}(r_{\min} + r_{\max})$ to preserve the symmetry of the distribution despite truncation. The function erf is the error function defined as

$$\mathrm{erf}(x) = \frac{2}{\pi} \int_0^x e^{-x^2}\, \mathrm{d}x. \tag{4}$$

Outside of $[r_{\min}, r_{\max}]$, we define

$$F_R(r) = 0, \qquad r < r_{\min}, \tag{5}$$

$$F_R(r) = 1, \qquad r > r_{\max}. \tag{6}$$

Hence, no cell divides up to a radius $r_{\min}$, whereas cells with a radius greater than $r_{\max}$ always divide. In a time discrete process, the division probability cannot be directly calculated from (3). Instead, previous attempts to divide have to be included as well. The exact division probability of a cell with radius $r^{(N)}$ at time step $N$ is then described by

$$p^{(N)} = \frac{F(r^{(N)}) - F(r^{(N-1)})}{1 - F(r^{(N-1)})}. \tag{7}$$

Following cell division, we use mass/volume conservation to calculate the daughter cell's radii $r_1$ and $r_2$ as

$$\frac{4}{3}\pi r^3 = \frac{4}{3}\pi r_1^3 + \frac{4}{3}\pi r_2^3. \tag{8}$$

In our model, we assume symmetric cell division, i.e. $r_1 = r_2 =: r_0$. This yields

$$r_0 = \frac{r}{2^{\frac{1}{3}}}. \tag{9}$$

### 2.3 Cell-cell adhesion and repulsion

The growth of a cell inevitably causes the displacement of itself and any adjacent cells. The corresponding equations of motion for $n$ cells were obtained by an overdamped approximation, i.e. inertia was neglected such that the velocities of the cells are proportional to the forces acting on them, i.e.

$$\frac{d\boldsymbol{x_i}}{dt} = F_0 \sum_{\substack{j=1 \\ j \neq i}}^{n} F_{i,j} \frac{\boldsymbol{x}_j - \boldsymbol{x}_i}{|\boldsymbol{x}_j - \boldsymbol{x}_i|}, \qquad \text{for } i = 1, ..., n. \tag{10}$$

Variable $\boldsymbol{x}_i$ describes the centroid of cell $i$. Parameter $F_0$ is a constant scaling factor, whereas $F_{i,j}$ describes the magnitude of the force of cell $j$ acting on cell $i$. The forces are derived from the Morse potential, which has already been successfully applied in similar biological contexts [8, 9]. They can be written in terms of cell radii $r_i$, $r_j$ and positions $\boldsymbol{x}_i$, $\boldsymbol{x}_j$ such that

$$F_{i,j} = \begin{cases} 2\alpha \left( e^{-\alpha(|\boldsymbol{x}_j - \boldsymbol{x}_i| - \sigma(r_j + r_i))} - e^{-2\alpha(|\boldsymbol{x}_j - \boldsymbol{x}_i| - \sigma(r_j + r_i))} \right) & \text{for } |\boldsymbol{x}_j - \boldsymbol{x}_i| \leq r_j + r_i, \\ 0 & \text{for } |\boldsymbol{x}_j - \boldsymbol{x}_i| > r_j + r_i. \end{cases} \tag{11}$$

The parameter $\alpha$ describes the stiffness of the cells, whereas $\sigma \in (0, 1]$ defines the optimal distance between two cells in contact as a fraction of the sum of their radii. The effect of the forces depends on the relative position of two cells. If the cells are too close, they will repel (Fig. II (a)), whereas they will adhere to each other if they are too far from each other but still in contact (fig. II (c)). In between, an optimal state will be found (Fig. II (b)). If the distance between the cells exceeds the sum of their radii, there can be no physical interaction (Fig. II (d)).

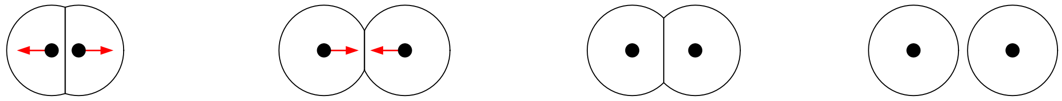

(a) Repulsion (b) Adhesion (c) Optimum (d) No interaction

Figure II: Illustration of the four different cases of interaction. Black dots represent the centroids of the cells. Black lines indicate their cell membranes. For repulsion (a) and adhesion (b), red arrows show the direction of the respective forces acting on each cell. For the optimal distance (c) as well as disconnected cells (d), no forces are present.

### 2.4 Transcriptional regulation

We employ our previously developed model for transcriptional regulation [10]. It follows the temporal evolution of two transcription factors $u$ and $v$ based on a model derived from statistical thermodynamics. The corresponding gene regulatory network (GRN) (Fig. III) involves mutual inhibition of $u$ and $v$ as well as their auto-activation. An external signal $s$ is used to inhibit $u$ while also activating $v$. For a tissue with $N$ cells, We end up with a system of ordinary differential equations (ODEs)

$$\begin{aligned} \frac{du_i}{dt} &= r_u \frac{\eta_u u_i}{1 + \eta_v v_i (1 + \eta_s \eta_{vs} s_i) + \eta_u u_i + \eta_s s_i} - \gamma_u u_i \\ \frac{dv_i}{dt} &= r_v \frac{\eta_v v_i (1 + \eta_s \eta_{vs} s_i)}{1 + \eta_v v_i (1 + \eta_s \eta_{vs} s_i) + \eta_u u_i + \eta_s s_i} - \gamma_v v_i, \qquad i = 1, ..., N, \end{aligned} \tag{12}$$

with the energy coefficients $\eta_x = e^{-\Delta \epsilon_x}$, the transcription rates $r_x$ and the decay rates $\gamma_x$.

We consider two types of external signal $s$. For the nearest neighbour signalling that is activated by $u$, we consider

$$s_i = \frac{1}{|N_G(i)|} \sum_{j \in N_G(i)} u_j. \tag{13}$$

Here, we used the notation $N_G(i)$ from graph theory to denote the neighbours of vertex $i$ in the graph $G$.

For the distance-based neighbour signal, $s$ is given as the result of the expressions of $u$ in any other cell according to

$$s_i = \left( \sum_{j \neq i} u_j q^{d_{ij} - 1} \right) \Big/ \left( \max_k \sum_{j \neq k} q^{d_{kj} - 1} \right), \qquad q \in [0, 1]. \tag{14}$$

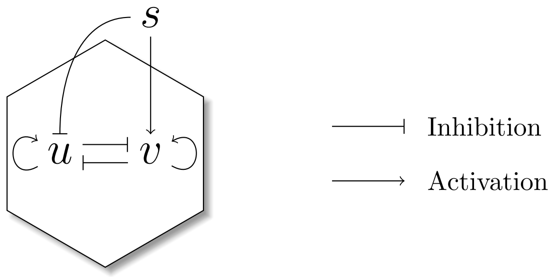


Figure III: Illustration of the gene regulatory network defining the temporal evolution of $u$ and $v$. Transcription factors $u$ and $v$ are regulated inside the cell while the signal $s$ provides extracellular input.

Here, $d_{ij}$ denotes the distance of cells $i$ and $j$ in the cell graph. By this definition of $s_i$, the parameter $q$ represents the strength of signal dispersion throughout the tissue. A value of $q = 0$ means that only direct neighbors influence a cell's fate. Increasing $q$ increases the influence of cells further away.

If transcriptional regulation is coupled with cell division, the values of $u_i$ and $v_i$ are evenly distributed between the daughter cells upon division.

### 2.5 Implementation

The implementation of the computational model is realized by a time discrete process. Beginning with a single cell, in each time step, the above mentioned steps are carried out sequentially. First the cell radii are updated via (2). Afterwards, the cell division probability is calculated according to (7). For each cell, we draw a random number from a uniform distribution between 0 and 1. If this number is less than the division probability, the cell is chosen for cell division. Thus, two cells are generated with equal radii according to (9). The axis of division is chosen randomly. The distance of the two cells after cell division is fixed as a parameter $h$. The amounts of $u$ and $v$ are equally distributed between the two daughter cells. Afterwards, the cell displacement is implemented by solving a single step of (10) numerically using the explicit Euler scheme. Finally, the transcriptional regulation is implemented by a second step of the explicit Euler scheme applied to the equations mentioned in [10]. The following tables further specifies the relevant parameters used in the simulation (Tables I and II). The evolution times were fitted to match average ICM cell counts of early (22), mid (25) and late (42) blastocysts.

| Model variable | Description |
|---|---|
| $u$ | Dimensionless concentrations of TF $U$ |
| $v$ | Dimensionless concentrations of TF $V$ |
| $s$ | Dimensionless concentrations of signal $S$ |

Table I: List of model variables and their descriptions.

| Model parameter | Fixed value | Description |
|---|---|---|
| $T$ | 108.95, 113.46, 131.74 | Evolution times for early, mid and late blastocysts |
| $N$ | 5000 | Number of time steps |
| $r^*$ | 1 | Maximum radius |
| $\lambda$ | 0.083 | Cell growth rate |
| $F_0$ | 0.01 | Displacement force scaling factor |
| $\alpha$ | 3 | Cell stiffness |
| $\sigma$ | 0.7 | Cell-cell distance optimality factor |
| $h$ | 0.23 | cell division distance |
| $-\Delta\varepsilon_u$ | 6-7.87 | Energy difference w.r.t. binding of $u$ |
| $-\Delta\varepsilon_v$ | 6 | Energy difference w.r.t. binding of $v$ |
| $-\Delta\varepsilon_s$ | 2 | Energy difference w.r.t. binding of $s$ |
| $-\Delta\varepsilon_u$ | 2 | Energy difference w.r.t. combined binding of $v$ and $s$ |
| $r_u$ | 1 | Transcription rate of $u$ |
| $r_v$ | 1 | Transcription rate of $v$ |
| $\gamma_u$ | 10 | Decay rate of $u$ |
| $\gamma_v$ | 10 | Decay rate of $v$ |
| $q$ | $\in (0,1)$ | Dispersion parameter for distance-based signal |
| $u_0$ | $\frac{3}{4}\frac{r_u}{\gamma_u}\left(1+\frac{\xi}{10}\right)$ | Initial condition for $u$ and $t=0$ |
| $v_0$ | $\frac{3}{4}\frac{r_v}{\gamma_v}\left(1+\frac{\xi}{10}\right)$ | Initial condition for $v$ and $t=0$ |
| $\xi$ | $\sim N(0,1)$ | Standard normal distribution noise |

Table II: List of model parameters and initial conditions together with their values and descriptions. Parameters downwards of $-\Delta\varepsilon_u$ are used in the transcriptional regulation model of [10].

[10] Simon Schardt and Sabine C. Fischer. Adjusting the range of cell-cell communication enables fine-tuning of cell fate patterns from checkerboard to engulfing. arXiv, 2022.
